# Interactions between the gut microbiome and mucosal immunoglobulins A, M and G in the developing infant gut

**DOI:** 10.1101/637959

**Authors:** Anders Janzon, Julia K. Goodrich, Omry Koren, the TEDDY Study Group, Jillian L. Waters, Ruth E. Ley

## Abstract

**Objective:** Interactions between the gut microbiome and immunoglobulin (Ig) A in infancy are important for future health. IgM and IgG are also present, however, their interactions with the microbiome in the developing infant are less understood.

**Design:** We employed stool samples sampled 15 times in infancy from 32 healthy subjects at 4 locations in 3 countries (from the TEDDY study). We characterized patterns of microbiome development in relation to levels of IgA, IgG and IgM. For 8 infants from a single location, we FACS-sorted microbial cells from stool by Ig status. We used 16S rRNA gene profiling on full and sorted microbiomes to assess patterns of antibody coating in relation to age and other factors.

**Results:** All antibodies decreased in concentration with age, but were augmented by breastmilk feeding regardless of infant age. Levels of IgA correlated with the relative abundances of OTUs belonging to the Bifidobacteria and Enterobacteriaceae, which dominated the early microbiome, and IgG levels correlated with *Haemophilus*. The diversity of Ig-coated microbiota was influenced by breastfeeding and age, but birth mode. IgA and IgM coated the same microbiota, while IgG targeted a different subset. *Blautia* generally evaded antibody coating, while members of the *Bifidobacteria* and Enterobacteriaceae were high in IgA/M.

**Conclusion:** IgA/M have similar dynamics with respect to microbiome development with age, and their interactions with the microbiome are influenced by breastfeeding status. IgG generally does not coat the commensal microbiota.

**Summary:** **What is already known on this subject?**

- Secretory IgA coats ~50% of microbiota in the gut
- IgM and IgG are less prevalent and coat a lower fraction in the adult, dynamics in the infant gut are not well characterized.
- Breastmilk is a source of IgA to the infant gut and decreases with time.
- IgA coating of microbial cells in infant gut microbiome decreases over time.

**What are the new findings?**

- Breastfeeding augments the IgA coating of the microbiome at all ages.
- IgA and IgM coat many of the same cells, whereas few are coated by IgG alone.
- Bifidobacteria, Enterobacteriaceae, *Ruminococcus gnavus* are enriched in IgA/M-coated cell fraction, *Blautia* is enriched in uncoated fraction.
- IgG levels correlated with *Haemophilus*.

**How might it impact on clinical practice in the foreseeable future?**

- Ig-coated fraction of the gut microbiome could serve as a useful tool for tracking development of the infant gut microbiome, and/or identifying aberrations to immune sensing of the microbiome.

## Introduction

The gut microbiota and the immune system and their interactions develop in tandem in infancy [1,2]. The IgA component of breast milk is protective against infection in immunologically immature infants, and may also direct the development of the gut microbiota. Immunoglobulin A (IgA) is secreted into the gut lumen where it binds with antigens from food and microbiota, thereby excluding them from direct contact with the host epithelial cells [3]. At birth, neonates generally have undetectable IgA in meconium [4], and it takes a few weeks for their immune systems to initiate the IgA production and secretion into the gut [5]. Breastmilk is an important early source of IgA and breastfeeding is associated with high levels of fecal IgA in infants [4,6]. Planer *et al* characterized the fraction of IgA-coated fecal microbiota in infants over the first few years of life and reported differences between breastmilk and formula fed infants, which may relate to differences in how the microbiota develop in these two groups [7].

Although IgA is the dominant antibody in the gut, IgM and IgG are also present. The adult gut sees up to 5 grams of secretory immunoglobulin A (IgA) daily, 100-fold less secretory IgM, and 1000-fold less IgG [8]. IgM and IgA are both produced by B cells locally, and the predominant class-switching that occurs in B cells of the gut-associated lymphoid tissue is from IgM to IgA. Both are secreted into the gut via the same mechanism (polymeric Ig receptor); IgM as a pentamer and IgA as a dimer. In contrast, IgG is the most common antibody in circulation, but can also be transported into the gut via a neonatal Fc receptor [9]. Whereas IgA/M are produced in response to luminal microbial epitopes that are sampled by dendritic cells, IgG induction is thought to require crossing of the barrier by antigens, such that IgG is not produced continuously in response to common gut antigens.

Based on its similarity to IgA, IgM may be expected to follow similar patterns of microbiota-binding, whereas IgG may not. In healthy adults, IgA has been shown to coat a greater proportion of the stool microbiota than IgG or IgM, but whether the diversity of taxa targeted by these antibodies differs has not been reported in controls [10]. To gain a baseline understanding of how IgA, IgG and IgM coat gut microbiota during microbiome development in infancy, here we performed a longitudinal analysis of the fecal microbiome of healthy infants in which we characterized the diversity of microbiota coated with IgA, IgM and IgG as a function of time and with respect to feeding regimen and antibody levels.

## Material and Methods

### Subjects and selection criteria

This study was conducted using fecal samples collected by the international type-1 diabetes (T1D) epidemiological prospective cohort study “The Environmental Determinants of Diabetes in the Young” (TEDDY) [11]. Subjects were screened for Type 1 Diabetes (T1D) risk through HLA genotyping, and all samples used in this study were collected from children with HLA genotypes that put them at high-risk of developing T1D. None of the subjects included in the current study have developed T1D or seroconverted for any of the 3 measured T1D-associated autoantibodies (glutamic acid decarboxylase autoantibodies, IA-2 autoantibodies, insulin autoantibodies) by 24 months of age and had not developed T1D by December 2016 (approximately 10 years of age). We focused on a set of longitudinal stool samples (9 to 16 timepoints per subject) obtained from 32 age and sex-matched healthy children (468 fecal samples) from the USA (Georgia and Washington), Germany, and Sweden. Subjects were excluded if they provided less than 12 longitudinal samples, if their participation and sample collection was through a long distance protocol, or if they dropped out of the study before or at 24 months of age. Additional data collected as part of the TEDDY study and that were used in the current analysis included physical descriptors (e.g., length, weight), diet, illnesses, hospitalizations, vaccinations and medicines, and social data such as daycare attendance.

### IgA, IgG, and IgM ELISAs

IgA, IgG, and IgM were quantified in duplicate for all samples. A serial dilution of reference human serum (Bethyl Laboratories, Montgomery, TX) was used for generating a standard curve, and blocking buffer was used as a negative control. The immunoglobulin concentrations were log transformed (with an offset of 0.01 added to IgM and IgG concentrations to handle zero values) before downstream analysis (see supplementary methods).

### FACS sorting of antibody-coated cells

Microbial FACS was performed on 117 fecal samples derived from 8 subjects from the Georgia study site. Samples were vortexed and centrifuged to separate bacterial cells from large particles and debris, and the resulting supernatant was transferred to a new tube. This supernatant was then resuspended in PBS and centrifuged to remove unbound immunoglobulins. The resulting pellet was then resuspended in PBS + 0.1% BSA before labeling with the respective IgA, IgG, and IgM fluorophore-labeled antibodies. Samples were incubated for 30 minutes and then washed twice prior to flow cytometry and cell sorting (see supplementary methods).

### Microbial diversity analysis

Genomic DNA was extracted from 468 fecal samples (approximately 20 mg per sample) using the PowerSoil - htp DNA isolation kit (MoBio Laboratories Ltd, Carlsbad, CA). The 16S rRNA V4 hypervariable region was then PCR amplified [12], and the resulting amplicons were pooled and sequenced using the Illumina MiSeq 2×250 bp paired end platform at Cornell Biotechnology Resource Center Genomics Facility.

Sequence analysis was performed using the open-source software package QIIME 1.8.0 (Quantitative Insights Into Microbial Ecology) [13]. Briefly, reads were quality filtered before using the open-reference OTU picking at 97% against the Greengenes August 2013 database. The data was rarefied at 18,429 sequences per sample, which was the lowest sample sequencing depth over 1,000, in order to include as many samples and sequences as possible (described in more detail in the online supplementary methods).

### Statistical analyses

#### Linear mixed models

All linear mixed effects models were fit using the lme4 [14] package in R. Significance was assessed using an F-test with a Satterthwaite approximation for denominator degrees of freedom calculated by the R package lmerTest. Post-hoc pairwise comparisons were performed using Tukey’s HSD tests implemented in the lsmeans [15] R package.

#### Association of subject characteristics with beta diversity and alpha diversity

A linear mixed model was used for associating individual metadata with alpha and beta diversity. The only factors used in the model examining the effect of time since exposure to oral antibiotics on unweighted UniFrac PC4 were: age, the time since oral antibiotic exposure, and the random effect for subject.

#### Correlations between immunoglobulin levels, age, and microbial diversity

The R package rmcorr [16] was used to calculate pairwise repeated measures correlations (and their significance) between levels of IgA, IgG and IgM, age, the first three unweighted UniFrac PCoA PCs, and Phylogenetic Diversity. A repeated measures correlation was used because it accounts for the multiple sampling of individuals.

For further details, see the online supplementary methods.

## Results

### The small sample set is representative of the larger TEDDY population

We looked for previously reported patterns of microbiota diversity in relation to metadata in order to establish that the small cohort used here is representative of the larger TEDDY cohort. Two recent papers have reported on the gut microbiome in the TEDDY cohort as well, employing a 16S rRNA survey on 903 subjects, and metagenomics with 783 subjects [17,18]. Here, we used a subset (32 subjects sampled longitudinally, unsorted stool) that were sequenced ~6X more deeply. Our results recapitulate those of Stewart *et al* and Vatanen *et al* in the following ways: (i) Bifidobacteriaceae and Enterobacteriaceae dominated the infant gut at earlier timepoints, while Firmicutes and Bacteroidetes increased in relative abundance later (see online supplementary figure S1); (ii) breastfeeding status was significantly associated with several PCs from both unweighted and weighted UniFrac PCoA (see online supplementary figures S2, S3; table S1) after correcting for multiple testing; (iii) the only other factor with a significant association to any PC was geographic location (see online supplementary figure S4A); (iv) we observed a weak association between antibiotic exposure and unweighted UniFrac PC4 (see online supplementary figure S4B-C); (v) age had a significant association with alpha-diversity (p<10^−10^, see online supplementary figure S5). Although these findings that age and breastfeeding status are associated with microbiome diversity are not novel, we report them here to underscore the independent repeatability of previous reports, and to establish that the subset of subjects that we used in subsequent analysis are representative of the larger TEDDY cohort.

### Stool antibody levels decrease with infant age and are related to breastfeeding status

We observed that levels of IgA, IgG and IgM (measured by ELISA in the same stool samples for which the microbiome was analyzed) were positively correlated to each other across the sample set (see online supplementary figure S6). Age (and therefore also microbiome alpha diversity) was negatively correlated with levels of IgA (repeated measures correlation *r*_*rm*_=−0.45, p=6.14 ⍰ 10^−22^), IgG (*r*_*rm*_=−0.37, p=1.13 ⍰ 10^−14^) and IgM (*r*_*rm*_=−0.23, p=3.29 ⍰ 10^−5^), although the IgG and IgM correlation with age was not as strong as that for IgA (see online supplementary figure S6). No association of geographic location or HLA genotype was observed with any of the immunoglobulin levels in stool.

We assessed the relationships between stool IgA levels, age and breastfeeding status to ask how IgA levels varied over time. We were particularly curious to see if the cessation of breastfeeding made an impact on stool IgA levels, and if the effect of breastfeeding was age-dependent. We found that IgA levels in stool decreased significantly with age (p=1.79 ⍰ 10^−11^; figure 1). Furthermore, we observed that breastfeeding was significantly associated with higher stool IgA levels (p = 0.039), and that this association was not dependent on the age of the infants. These observations corroborate previous reports that IgA levels decrease with age, but that at any age, breastmilk delivers additional IgA to the infant.

**Figure 1.**
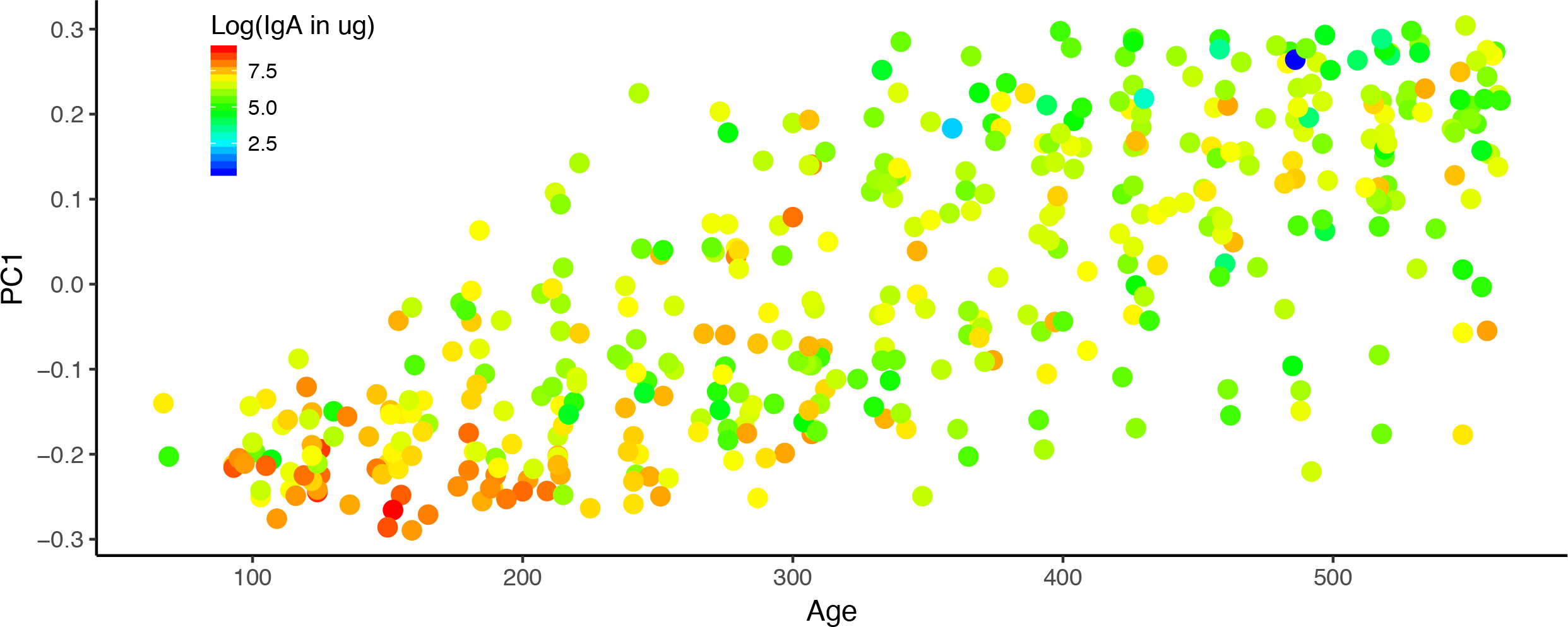
Infant fecal IgA concentrations over the first couple years of life. PC1 from the PCoA of the fecal microbiome unweighted UniFrac distances is plotted against the age of the infant in days at the time of sampling. The points are colored by the log transformed fecal IgA concentrations; lower concentrations are blue and higher concentrations are red.

Next, we examined the association between stool levels of IgA, IgG, and common OTUs (e.g., those shared by greater than 40% of samples; figure 2). IgA levels were positively associated with several Enterobacteriaceae OTUs and a few Bifidobacteriaceae OTUs. Only one OTU was associated with IgG levels in feces: this OTU belonged to genus *Haemophilus* (BH adjusted p=1.66 ⍰ 10^−3^).

**Figure 2.**
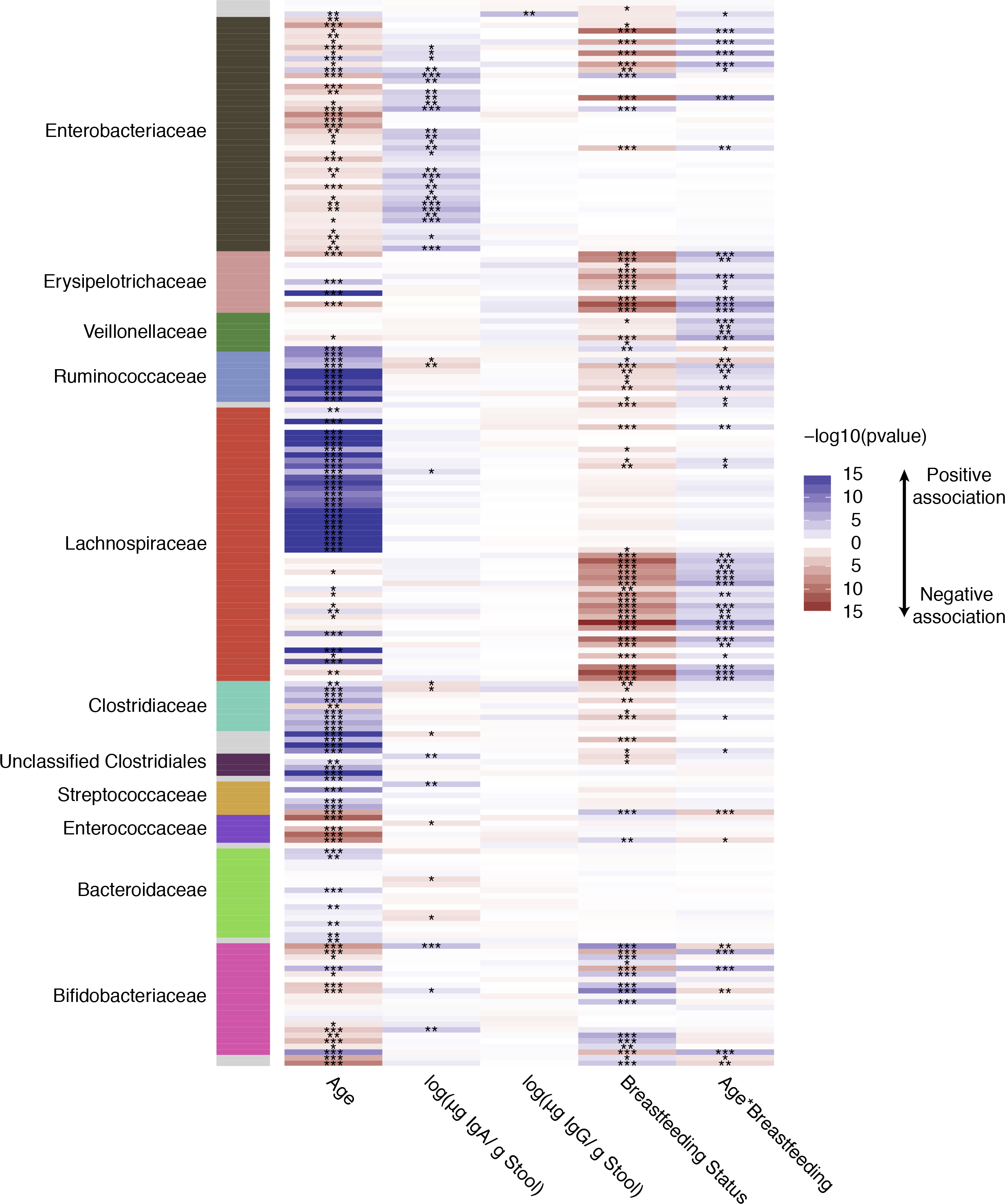
A heatmap summarizing the association of age, fecal IgA and IgG concentrations, and breastfeeding status with common OTUs in the infant fecal microbiomes. Along the vertical axis are each of the common OTUs (non zero value in >40% of samples tested) and the horizontal axis is each of the fixed factors examined. The heatmap is colored by the −log10 of the p value from a linear mixed model examining the OTU association with the fixed effects; Blue indicates a positive association and red a negative association while white is a p value of 1, and the darker the color the more significant the association. The panel on the left is colored by the family level taxonomic association of the OTU.

### IgA and IgM coat many of the same microbial cells

For the 8 infants from Georgia, for whom 15 timepoints each were available, we sorted cells according to their antibody-coating status using FACS, using their distributions in four quadrants (see supplementary methods). The gatings for these four populations and the resulting quadrants are illustrated in figure 3B. We ensured that cross-reactivity of the anti-IgA, IgM and IgG antibodies was minimal, thus the patterns observed by flow cytometry indicate that many microbial cells were tagged with multiple antibodies. The gating patterns indicate that IgA and IgM coat many of the same cells, since the patterns of coating overlap. IgG-coated cells, on the other hand, fell into three categories: Q1, those low in IgG and high in IgA/IgM; Q2, those highly coated in IgG, IgA and IgM; Q3, uncoated cells; and Q4, those highly coated in IgG and low in IgA/IgM. We observed that the correlation between IgA and IgM coating was consistently higher than the correlation between IgA and IgG coating (p<10^−10^; figure 3C)

**Figure 3.**
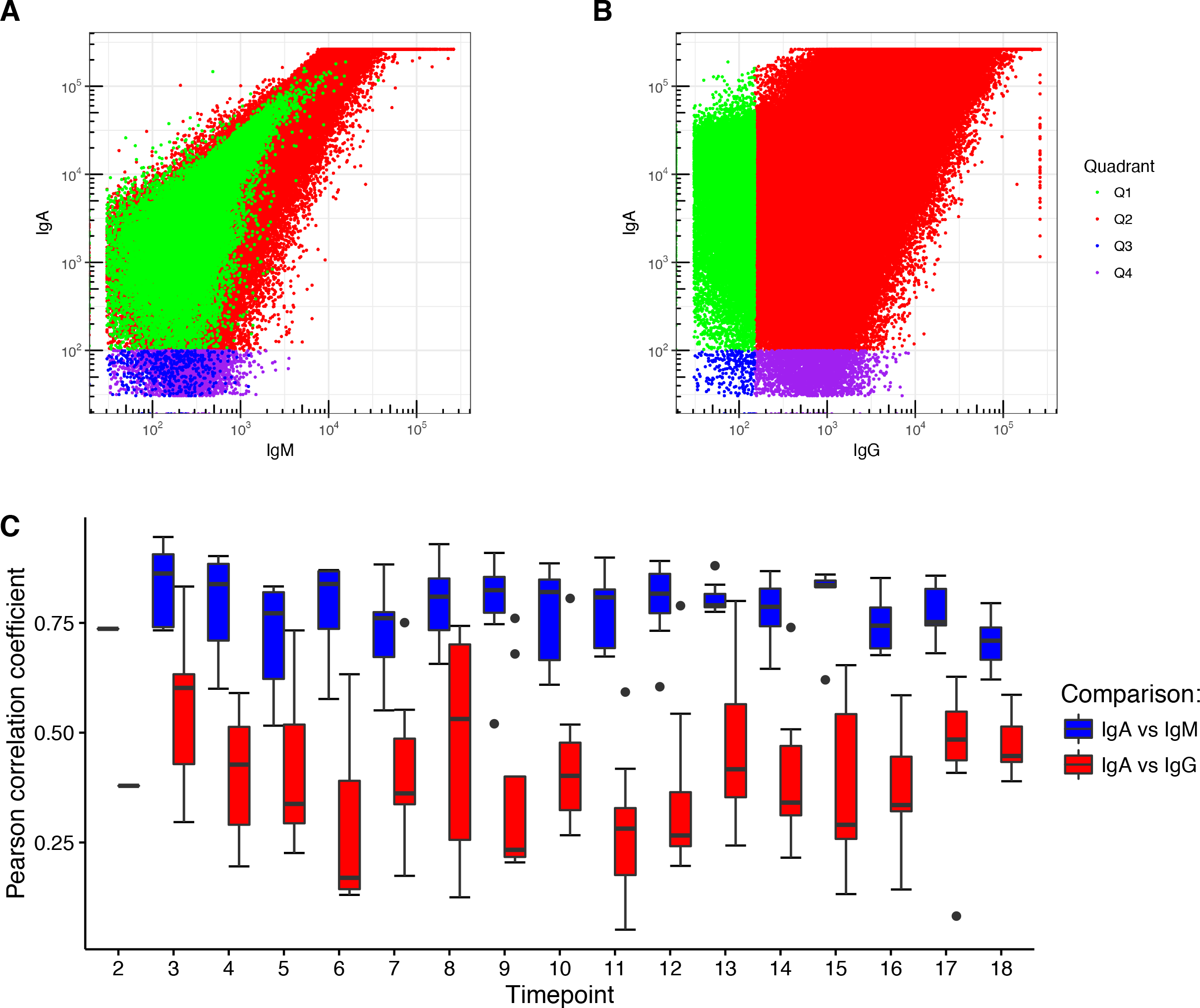
IgA and IgM coat many of the same microbial cells. (A) Representative data for FACS sorting of microbial cells in infant fecal samples. Frequencies of IgM versus IgA tagged cells (green and red, respectively). (B) This panel shows how the four Quadrants were gated: Q1-Q4 are indicated on the figure. Q1: IgA, IgM both high; Q2: All three high; Q3: all three low; Q4: IgG high, others low. (C) Box-plots at each time point of the Pearson correlations (y-axis) between IgA and IgM signals and between IgA and IgG signals from the FACS sorting of each fecal sample.

### Specific taxa discriminate total and FACS-sorted populations

We compared the diversity of the sorted microbiota (all Qs combined) to the whole microbiota (unsorted) to assess the impact of the FACS process on microbial diversity. We observed that the microbiota composition of the sorted cells (all Qs) is distinct from the total microbiome: a combined PCoA analysis of unweighted UniFrac analysis shows clear separation of unsorted and sorted cells along PC2 (figure 4A). The unsorted population was richer (Chao 1, Phylogenetic Diversity, Observed Species) and exhibited greater evenness (Gini coefficient) than all sorted cells (p<10^−10^ for all metrics; see online supplementary figure S7). We applied linear mixed models restricted to identify taxa that were differentially abundant between the unsorted and sorted populations (all Qs). This analysis identified members of the Bacteroidetes, Verrucomicrobia, gamma-Proteobacteria and most Firmicutes as comparatively enriched in the unsorted fraction, and Actinobacteria and alpha-Proteobacteria as enriched in the sorted fraction (figure 4C, see online supplementary figure S8). The difference in composition between sorted and unsorted fractions likely stems from the FACS process itself: cells belonging to Actinobacteria and alpha-Proteobacteria may be less likely to clump than others. Thus, the difference between sorted and unsorted microbiomes introduces the caveat that subsequent analyses with the sorted data are performed on a subset of the microbiome.

**Figure 4.**
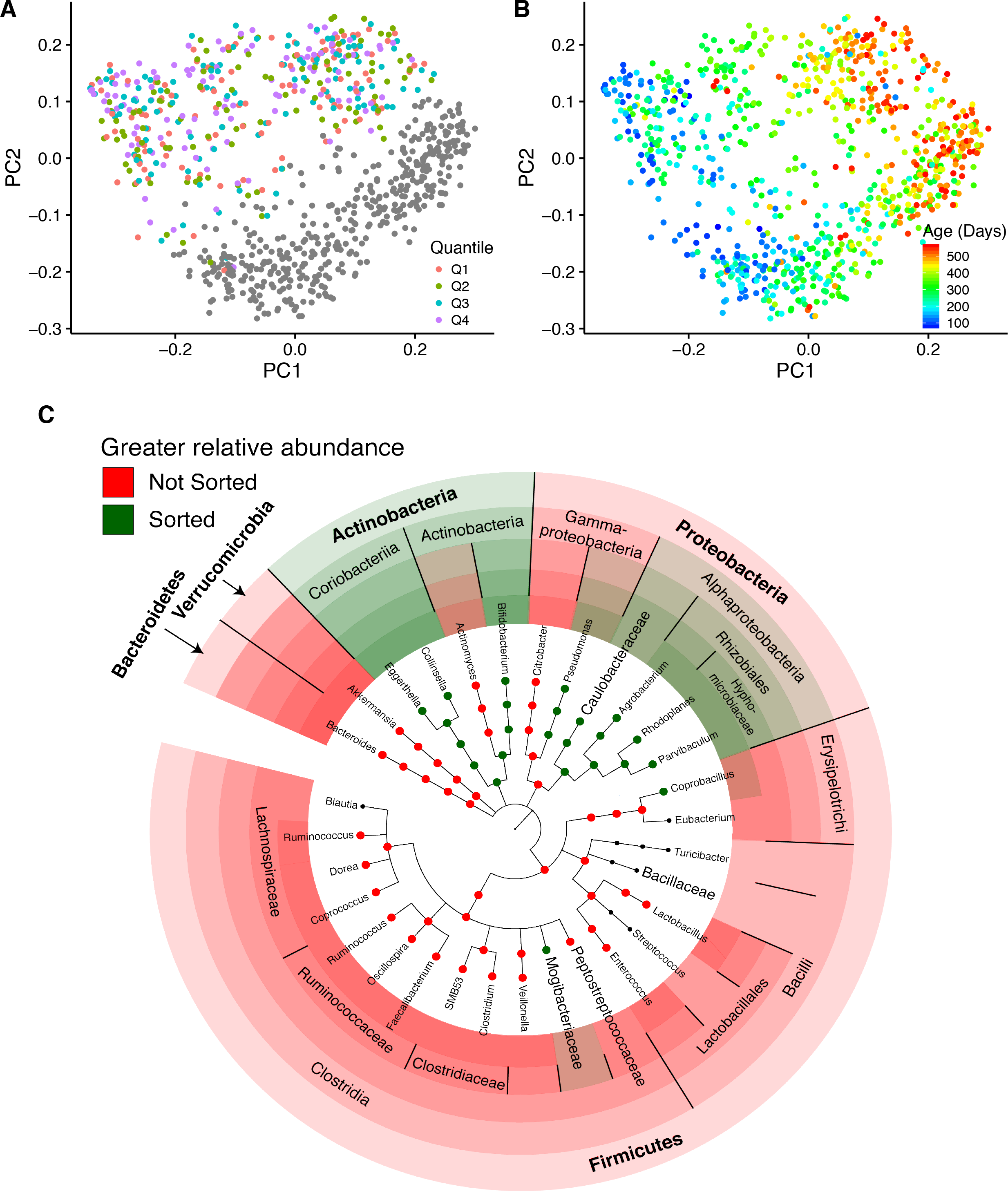
Fecal microbiome composition is significantly altered by FACS, but remains significantly associated with infant age. (A) PCoA of unweighted UniFrac distances colored by FACS quadrant. Unsorted samples are shown in gray. (B) PCoA of unweighted UniFrac distances colored by participant age. (C) Summary of taxa that are differentially abundant between the total (red) and FACS (green) populations.

### Antibody-targeted cells exhibit patterns similar to those of the total population

As observed for unsorted cells, the majority of the variation in the sorted cells (PC1 of the unweighted UniFrac PCoA) was explained by age (figure 4B, see online supplementary figure S9). Similarly, as age increased, alpha-diversity also increased (see online supplementary figure S7). Analysis of variance indicated that FACS quadrant was significantly associated with most of the first 10 PCs from the PCoA (restricted to only sorted 16S data) of both unweighted and weighted UniFrac and all four alpha-diversity metrics. Post-hoc pairwise comparisons showed that this association was driven primarily by a difference between Q4 (high IgG-only) and the other three quadrants. This finding might represent an overall diversity difference between the high IgG-only cells compared to others. However, the number of cells sorted into the high-IgG (Q4) quadrant was significantly lower than the other three quadrants (see online supplementary figure S10). Indeed, most of the cells coated in IgG were also coated by IgA and IgM, and are therefore sorted into Q2 rather than Q4. This significant difference in cell number between the quadrants could explain the difference in diversity observed between Q4 and the other three quadrants.

After exclusion of Q4 (high-IgG only), we observed some significant associations between the first 10 PCs of the beta diversity PCoA and the FACS quadrant, as well as several associations with infant age and infant breastfeeding status. The significant associations with unweighted UniFrac were with age (PCs 1, 2, and 3), breastfeeding status (PCs 1, 2, 3, 6, and 7), and FACS quadrant (PCs 7 and 10; see online supplementary table S2). Further analysis of the FACS quadrant associations showed that Q3 (uncoated) was different from both Q1 (p = 0.030) and Q2 (p = 0.001) along PC7, and that Q2 differed from Q3 (p = 0.010) along PC10 (post-hoc pairwise comparisons between the quadrants using Tukey’s HSD method to adjust for multiple comparisons). Among the weighted UniFrac PCs, many are associated with FACS quadrant (PCs 1, 5, 7, 8, and 9; post-hoc analysis shows that Q3 is different from Q1 and Q2), PC1 is associated with breastfeeding status, and PC2 with age (see online supplementary table S2). These results indicate that the diversity of cells targeted by antibodies is influenced by breastfeeding status and with infant age. Furthermore, the specific combination of antibodies on the cells is not random.

### Variability in the relative abundance of specific taxa between FACS quadrants

We next identified specific OTUs differentially abundant between the quadrants. To identify common OTUs (i.e., OTUs with non-zero sequence counts in greater than 40% of FACS samples examined) with differential relative abundance between quadrants, we performed linear mixed models using each OTU as a response variable. We searched for differences between the coated (Q1 and Q2; excluding Q4 because of low cell population) and uncoated (Q3) populations. When comparing Q1 to Q3, we found significant differences in the relative abundances of 80 out of 190 OTUs (figure 5, see online supplementary figure S11 and supplementary table S3). These included mostly OTUs classified as *Blautia*, which had higher relative abundance in Q3 (uncoated) compared to other Qs. OTUs that were higher in Q1 (IgM and IgA both high) were mostly classified as *Bifidobacterium*, unclassified Enterobacteriaceae, and *Ruminococcus gnavus*. Similarly, 101 OTUs had differential relative abundances between Q2 (all high) and Q3 (all low), with *Blautia* significantly higher in Q3 (see online supplementary figure S12 and supplementary table S4).

**Figure 5.**
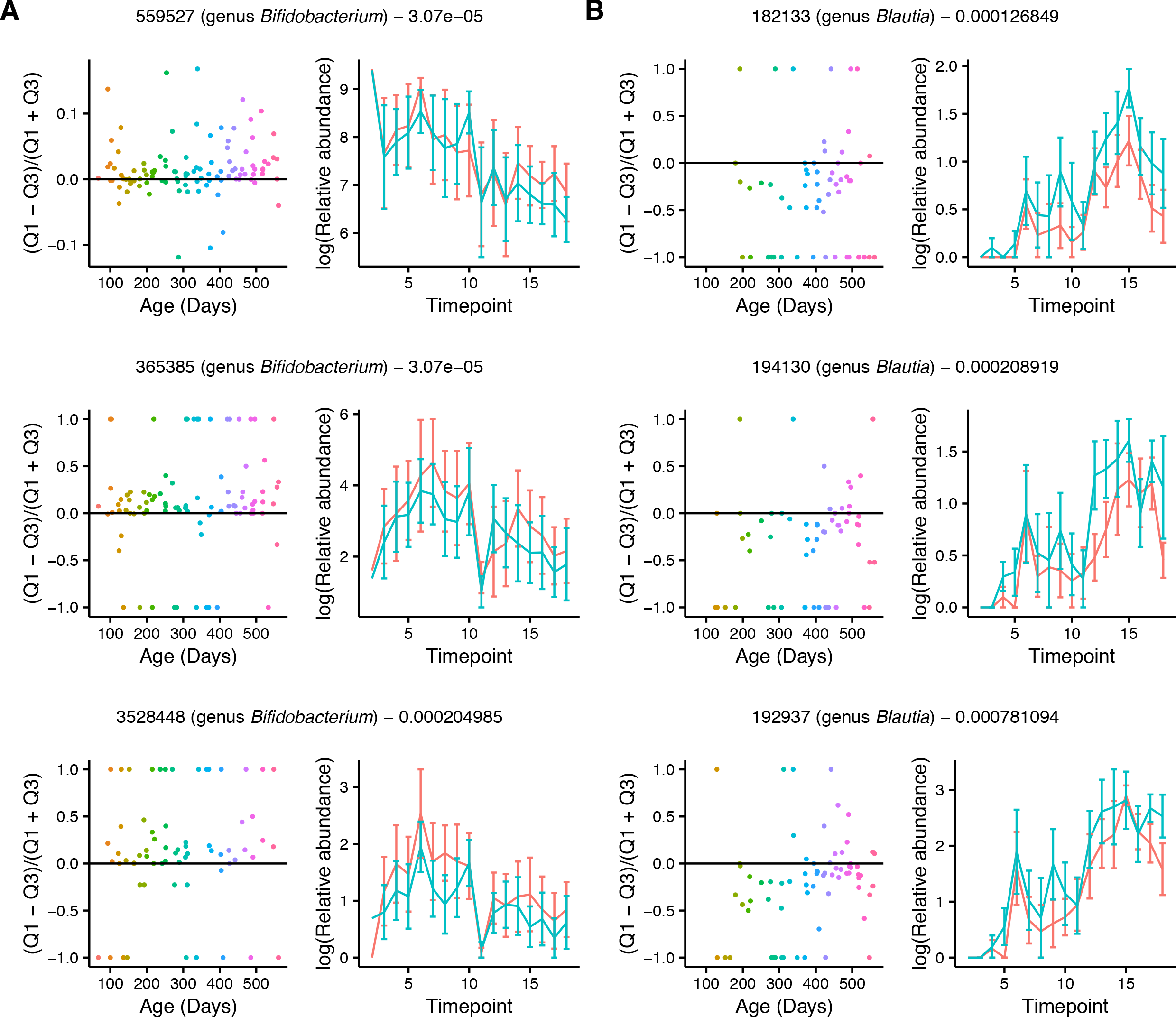
Many differentially abundant OTUs between Q1 (IgM and IgA both high) and Q3 (low coating) are classified as *Blautia* and *Bifidobacterium*. (A-B) Plots for a subset of differentially abundant OTUs between Q1 and Q3 illustrating (A) higher relative abundance of *Bifidobacterium* OTUs in Q1 and (B) higher relative abundance of *Blautia* OTUs in Q3 (see online supplementary figure S11 for plots of all OTUs). Plots on the left show the log(Q1 OTU abundance) − log(Q3 OTU abundance) divided by log(Q1 OTU abundance) + log(Q3 OTU abundance) over time where each point represents a sample and is colored by timepoint. Positive values on the y-axis indicate enrichment of the OTU abundance in Q1 and negative values indicate and enrichment of the OTU abundance in Q3; this is similar to the IgA index defined in Planer *et al* 2016. Plots on the right show the average of the log transformed OTU relative abundance in Q1 (Red) and Q3 (Blue) at each time point. OTU Greengenes ID, taxonomic classification, and the p value from the linear mixed model using OTU as a response variable is indicated above each set of graphs. Only common OTUs (non-zero value in >40% of samples tested) were tested.

### Cross-validation for IgA-targeted microbiota

We compared the results of our analysis with those of Planer *et al*, who used FACS followed by 16S rRNA gene sequencing to characterize the IgA coated microbes of 40 twin pairs over the first couple years of life [7]. We used the same reference database as Planer *et al* to classify sequences, therefore we could compare OTUs directly. Only two of the OTUs that the Planer study identified as differentially abundant between the IgA^+^ and IgA^−^ fractions were also among the common OTUs examined in our analysis, but interestingly, both behaved similarly with respect to antibody coating. Greengenes OTU 365385 (genus *Bifidobacterium*) was consistently targeted by IgA in the Planer study and we observed that it was proportionally higher in both Q1 (BH adjusted p = 3.07 × 10^−05^; figure 5A) and Q2 (BH adjusted p = 1.88 × 10^−05^) compared to Q3. The other Greengenes OTU detected in both datasets (606927; family Peptostreptococcaceae) was consistently non-targeted by IgA in the Planer study, and similarly, we observed this OTU to have higher relative abundance of this OTU in Q3 (uncoated) compared to Q2 (BH adjusted p = 0.013) (see online supplementary figure S12 supplementary table S4). These comparisons indicate that taxon-specific antibody coating profiles can be generalizable across studies.

## Discussion

This study included infants from four distinct geographic locations, two in the USA and two in Europe. These geographic locations left a discernible imprint on the infant microbiome, which was observed in other studies of the microbiome in the TEDDY group [17,18]. Effects of breastfeeding status on the microbiome and antibody levels, and the decrease in antibody levels with age were also similar across subjects, regardless of the shift in diversity associated with the geographic location of the infants. We also report a strong impact of breastfeeding status and age on gut microbial community structure, which is consistent with the two other TEDDY studies [17,18]. In contrast to these larger studies, we did not observe an effect of birth mode on microbiome diversity in this dataset.

Our longitudinal analysis of the Georgia USA infants’ gut microbiota targeted by antibodies over time revealed that IgM and IgA target the same microbial populations. IgA and IgM targeting of the same cell population implies they may be induced via the same mechanism, and that like IgA, IgM induction is a local phenomenon. In contrast, we observed patterns of IgG coating were quite distinct from those of IgA and IgM. IgG is induced as a result of the systemic immune system’s recognition of a “non-self” antigen and is not, in general, produced locally in the gut. Thus, whereas IgA/M responses mirror the gut microbiome generally, the bacterial targets of IgG are related to those targeted by the systemic immune system around the time of collection. We noted a fair amount of overlap between the IgA/M and IgG coated fractions, however, suggesting redundancy across all three classes of antibody.

The majority of IgA secreted to the gut is polyclonal and thought to be relatively unselective as it binds with epitopes that are widely shared amongst gut bacteria [19,20]. In addition, secretory IgA affinity maturation by somatic hypermutation is a prominent feature of the human repertoire [21] and a large fraction of secretory IgA may be targeted to specific bacterial species [22]. Hapfelmeier and colleagues showed with a reversible colonization system in germfree mice that a specific strain of *E. coli* induced a sustained IgA response even after the strain was no longer there (>100 days), but that colonization with other bacteria induced a decline in the *E.coli*-specific IgA and an increase in IgA response to the newly introduced bacteria [23]. These results suggested the presence of a long-lived compartment of IgA secreting plasma cells in the intestinal *lamina propria*, but that the compartment has a finite size, so that the IgA secreted into the intestine depends on the dominant luminal microbial species. Based on this model, we would expect the IgA-coated fraction of microbiota to mirror the unsorted microbiota, and allowing for the caveats of FACS sorting (e.g., some phyla aren’t well represented in the sorted cells for reasons that are not well understood) this is the general pattern that we observed.

A few OTUs provided exceptions to the general pattern in that some OTUs were more highly represented in the IgA-coated than uncoated cell populations. In particular, OTUs classified as *Bifidobacterium*, unclassified Enterobacteriaceae, and *Ruminococcus gnavus* had higher representation in the IgA-coated cell fraction. In addition, we observed that levels of IgA antibody in stool was correlated to a few Bifidobacteriaceae OTUs and several Enterobacteriaceae OTUs. Planer *et al* also observed one of the same OTUs as highly coated in IgA in the fecal samples of children from Malawi [7], and Bifidobacteria in particular have been shown to induce high levels of IgA in the gut [24]. Bifidobacteria and Enterobacteriaceae dominated the infant gut microbiomes early on, however the higher-than expected levels of specific OTUs belonging to these taxa in the IgA-coated fraction suggests they may be stimulating IgA production specific to them in excess of what is expected from their relative abundances in the microbiota. Alternatively, given that here we characterize antibody coating of bacteria in the feces, these taxa may be preferentially cleared from the intestine once coated with IgA.

In contrast to the high-IgA-coated microbiota, OTUs classified as *Blautia* showed a lack of antibody coating. In the healthy mouse cecum, specific OTUs have been shown to be less coated than what would be expected by chance [25]. One explanation could be antibody evasion by regulation of epitopes in response to the antibodies. Certain bacterial species, including the gut commensal *Bacteroides thetaiotaomicron*, have been shown to downregulate the expression of epitopes in response to IgA [26–28]. Alternatively, the relative undercoating of specific taxa in stool may reflect patterns of clearance, and here uncoated *Blautia* OTUs are more likely to be shed than their coated counterparts. How IgA coating relates to the microbial ecology of the microbiota, their growth rates, the immune response to specific epitopes, and ultimately shedding of the cells, is complex and not well understood.

Although a large proportion of sorted cells was coated by all three antibodies, we did observe differences between the patterns of IgG coating compared to the patterns of IgA/IgM coating, although the low cell count in the IgG-only coated category stymied the interpretation. However, one OTU was associated with IgG levels in feces: this OTU belonged to the genus *Haemophilus*. IgG is produced systemically in response to infection, and it is also produced in response to immunization. Thus, this pattern may have resulted from the *Haemophilus* vaccine, which is given to infants. It is one of the few vaccines against a bacterium, and it is not a mucosal vaccine. *Haemophilus* colonizes the upper respiratory mucosa and not all are pathogenic. The *Haemophilus* that we observed here may include commensal specie(s) that share capsule epitopes with the vaccine strains.

Our results indicate similar dynamics of antibody levels and microbiome development with age and breastfeeding status in infants from different locations. Patterns of IgA/M patterning indicate that sorting for IgM coated cells may not be any more informative than sorting for IgA alone. IgG-coating of the infant gut microbiome may reveal antigen exposure if gut microbes cross-react. These data provide a baseline reference for further investigation of healthy or unhealthy children’s gut microbiomes in infancy.

**Table 1**. Summary of participant characteristics.

## Acknowledgements

We thank William Melvin and Wei Zhang at Cornell University for their assistance. Special thanks to Timothy Bushnell, Matthew Cochran and staff at the Flow Cytometry Core at the University of Rochester Medical Center. This work was supported by a National Science Foundation Graduate Fellowship (JKG), Swedish Research Council grant 2011-922 (AJ), and the Max Planck Society (REL). The TEDDY study is funded by U01 DK63829, U01 DK63861, U01 DK63821, U01 DK63865, U01 DK63863, U01 DK63836, U01 DK63790, UC4 DK63829, UC4 DK63861, UC4 DK63821, UC4 DK63865, UC4 DK63863, UC4 DK63836, UC4 DK95300, UC4 DK100238, UC4 DK106955, and Contract No. HHSN267200700014C from the National Institute of Diabetes and Digestive and Kidney Diseases (NIDDK), National Institute of Allergy and Infectious Diseases (NIAID), National Institute of Child Health and Human Development (NICHD), National Institute of Environmental Health Sciences (NIEHS), Juvenile Diabetes Research Foundation (JDRF), and Centers for Disease Control and Prevention (CDC). This work supported in part by the NIH/NCATS Clinical and Translational Science Awards to the University of Florida (UL1 TR000064) and the University of Colorado (UL1 TR001082).

