## Supplementary materials for "Interactions between the gut microbiome and mucosal immunoglobulins A, M and G in the developing infant gut"

**Supplementary Methods**

**Sample collection and treatment -** The samples used in this study were part of an existing sample collection [[1]](https://paperpile.com/c/IdncC4/bTLUc). In brief, fecal samples had been collected either directly at one of the TEDDY associated primary care centers in Germany, Sweden and the USA (Georgia and Washington) or collected at home and transported to the primary care facilities. Samples were then stored at -20 °C before transport on dry ice to the TEDDY clinical center at the University of Tampere, Tampere, Finland where they were diluted to a 1:10 suspension in Hank’s Balanced Salt Solution (HBSS), aliquoted and stored at -80°C. Aliquots of 1.0-1.5 mL were then shipped on dry ice to Cornell University (NY, USA), where they were stored at -80°C. For the subsequent steps of ELISA, FACS and fecal total DNA isolation, samples were always handled inside a laminar flow hood, with prior UV-treatment of all instruments and laboratory consumables, to prevent contamination. Due to the minimal starting material (≤1,000,000 cells) present in bacterial FACS samples, DNA isolation from these samples was performed in a clean lab such that extraction blank 16S rRNA PCR reactions yielded no bands at 40 cycles of amplification.

**IgA, IgG, and IgM ELISAs**

To prepare samples for ELISA quantification of IgA, IgG and IgM, aliquots of 468 samples from the 32 selected subjects were thawed and briefly vortexed to resuspend contents. 150 µL of each sample was then vigorously vortexed followed by centrifugation at 8,000 g for 10 minutes to pellet the solids. Pellets were stored at -80°C and used in subsequent extraction of fecal total DNA, and supernatants were immediately diluted 2-fold, 25-fold and 2000-fold in 1X PBS at pH 8.0 with 1% BSA (essentially globulin free) and 0.05% Tween 20 and used for ELISAs of IgG, IgM and IgA, respectively. All incubations were performed at ambient temperature unless specified, and plates were washed 5 times between all steps using an excess of PBS with 0.05% Tween 20.

Briefly, medium binding flat bottom 96-well plates (Santa Cruz Biotechnology, Dallas, TX) were coated for 1 hour with an Anti-Human IgA/IgG/IgM (H+L) polyclonal antibody (Sigma Aldrich) diluted to 10 µg/mL in 0.05 M Carbonate-Bicarbonate buffer at pH 9.6. Plates were then incubated overnight at 4 °C using a blocking solution of 1% BSA in PBS. Samples were added in duplicate to 96-well plates together with a dilution series of human reference serum (Bethyl Laboratories, Montgomery, TX) as a standard curve, or blocking solution alone as a negative control, and plates were incubated for 1 hour. Horseradish peroxidase-conjugated detection antibodies specific for human IgA, IgG or IgM (Bethyl Laboratories) were added at final concentrations of 11.1 ng/mL, 13.3 ng/mL and 15.4 ng/mL, respectively, and incubated for 1 hour. TMB enzyme substrate (Sigma, St. Louis, MO) was then added and the ELISA was developed for 15 min in the dark. The reaction was stopped by addition of 0.18 H_2_SO_4_ and absorbance was measured at 450 nm in a Synergy H1 plate reader (BioTek, Winooski, VT). Sample concentrations were calculated using the instrument software and 4-parameter curve fitting. The immunoglobulin concentrations were log transformed (with an offset of 0.01 added to IgM and IgG concentrations to handle zero values) before downstream analyses.

**FACS sorting of antibody-coated cells**

A total of 117 fecal samples derived from 8 subjects from the Georgia study site were selected for further analysis by microbial FACS. To prepare samples for FACS, the fecal suspensions were thawed and vigorously vortexed before 600 µL was transferred to a new microfuge tube. The tubes were centrifuged at low speed (15 g for 10 minutes) to separate bacterial cells from larger particles, and 450 µL of the supernatant was transferred to a fresh tube. Another 75 µL of the supernatant of all samples processed each day was mixed in a single tube to provide a non-stained control of background fluorescence. In addition, an aliquot of a premixed suspension of 4 samples not included in our study was thawed and used to generate 4 single-stained controls for instrument calibration.

PBS (1 mL) was added to each tube, vortexed briefly and then centrifuged at 8000 g for 10 minutes. The supernatant was then carefully aspirated and decanted to remove non-bound immunoglobulins. The pellet was then resuspended in 450 µL PBS with 1 % BSA and 10 µg/mL each of fluorophore-labeled polyclonal goat F(ab)_2_ anti-human IgA, IgG and IgM antibodies. The antibodies used were Alexa 488-conjugated anti-human IgA (Life Technologies, Carlsbad, CA, USA), APC-conjugated anti-human IgG (Jackson Immunoresearch Laboratories, West Grove, PA, USA) and R-PE-conjugated anti-human IgM (Jackson Immunoresearch Laboratories). Aliquots for fluorophore-antibody single-stained controls were resuspended in PBS/BSA with only a single antibody at identical concentration; non-stained controls were incubated with PBS/BSA alone. Samples and controls were then incubated for 30 minutes at room temperature, followed by two wash steps with 1 mL PBS and centrifugation at 8,000 g for 10 minutes. Finally, each sample pellet, and one control pellet, was resuspended in 450 µL PBS with 1:500 dilution of Vybrant DyeCycle Violet as a cell stain. Remaining single-stain controls and the non-stained control were resuspended in PBS alone. The final suspensions were stored on ice until FACS analysis, between 2 and 8 hours.

Flow cytometry and cell sorting was performed at the University of Rochester Medical Center Flow Cytometry Core Facility on a FACSAria II (BD Biosciences, San Jose, CA, USA) equipped with 4 lasers: Violet 407 nm (100 mW), Blue 488 nm (100 mW), Green 532 nm (150 mW) and Red 633 nm (40 mW). The instrument was operated with the following laser, filter and fluorophore combinations: 407 nm laser with BP at 450/50 for Violet DyeCycle (cell marker), 488 nm laser with LP at 505 and BP at 525/50 for Alexa 488 (IgA), 532 laser with BP at 575/25 for R-PE (IgM) and the 633 nm laser with BP at 660/20 for APC (IgG). The 488 nm laser was also used to collect FSC and SSC (with BP at 488/10). Pulse geometry gates in FSC and SSC spectra were used to exclude cell multiplets and a threshold was set for Violet DyeCycle fluorescence to exclude non-cellular particles. Since initial experiments showed strong correlation between IgA and IgM signals, the instrument was then set up for 4-way sorting from quadrants (Q1-4) in the IgA vs IgG dotblot, allowing collection of cells with the following profiles: Q1: IgA+IgM both high and IgG low; Q2: All high; Q3: All Low; Q4: IgG high, IgA and IgM both low.

The instrument was calibrated at the start of each day using the single- and non-stained controls. For each sample, 100,000 events were collected to generate cytometry data and then sorted a minimum of 500,000 cells from each of Q1-3 and a minimum of 50,000 cells from Q4. The FACS instrument was purged for 3 minutes between each sample, to ensure that cells and debris were flushed through completely and to allow event counts to return to background levels. Cells were sorted into 13 mL Falcon tubes. Immediately following cell sorting, volumes corresponding to 500,000 - 1,000,000 cells (Q1-3) or 50,000-500,000 cells (Q4) were transferred to 2 mL microfuge tubes and centrifuged at 18,000 g for 30 minutes at 4 °C to collect the sorted cells. The supernatant was carefully aspirated and tubes with pellets were stored at -20 °C until transportation on dry ice to Cornell University, Ithaca, where they were stored at -80 °C until DNA extraction. Before usage, all tubes had been UV-treated twice for 15 minutes in a biosafety hood to minimize background DNA.

**DNA extraction, 16S rRNA gene PCR and sequencing**

Genomic DNA was isolated from approximately 20 mg of pelleted material from 468 fecal sample pellets using the PowerSoil - htp DNA isolation kit (MoBio Laboratories Ltd, Carlsbad, CA). DNA-free water, as a negative extraction control, was added instead of sample to at least 10 wells in each 96-well plate. To maximize DNA recovery from the 468 samples of sorted cells, total DNA was isolated using a simple alkali-heat treatment directly in the microfuge tubes. Briefly, 50 µL of 0.2 N KOH was added to each tube, followed by 60 minutes incubation in a heating block at 98 °C. After heat treatment, 25 µL 0.4 M Tris*HCl (pH 7.5) was added. Finally, 25 µL 0.4 N HCl was added.

Amplification and sequencing of the V4 hypervariable region of the 16S rRNA gene from the isolated DNA was performed as described previously [[2]](https://paperpile.com/c/IdncC4/ykue2), with modifications for the DNA recovered from sorted cells to optimize PCR efficiency for low DNA concentration. For the fecal pellet DNA, the PCR reaction mixture consisted of 2.5 U Easy-A high-fidelity polymerase, 1 × buffer (Stratagene, La Jolla, CA), 100 ng DNA template, and 0.05 µM of each primer (515F and 806R) and the reaction conditions were an initial denaturation step at 94 **°**C for 3 minutes followed by 25 cycles of denaturation at 94 **°**C for 45 seconds, annealing at 50 **°**C for 60 seconds, extension at 72 **°**C for 90 seconds, and a final extension at 72 **°**C for 10 minutes. For DNA from sorted cells, polymerase and buffer concentrations were the same, but 10% (V/V) DNA template and 0.1 µM of each primer were used. PCR conditions were kept the same, but the reaction was allowed to continue for 35 cycles. PCR reactions for all samples, fecal pellets and sorted cells, were carried out in duplicate 96 well plates, with one aliquot reaction per plate. Apart from the negative extraction controls described above, an additional 4-6 negative PCR controls were included per plate. Replicate PCR reactions were combined and purified using a magnetic bead system (Mag-Bind® EZPure, Omega Bio-Tek, Norcross, GA). PCR amplicons were quantified using the QuantiT PicoGreen dsDNA Assay Kit (Invitrogen, Carlsbad, CA). Aliquots of amplicons (at equal masses) were combined for a final concentration of approximately 15 ng/µl. DNA was sequenced using the Illumina MiSeq 2x250 bp paired end platform at Cornell Biotechnology Resource Center Genomics Facility.

**Microbial diversity analysis**

The sequences were analyzed using the open-source software package QIIME 1.8.0 (Quantitative Insights Into Microbial Ecology) [[3]](https://paperpile.com/c/IdncC4/wMpP3). Matching paired-end sequences (mate-pairs) were merged using fastq-join in the ea-utils software package with a minimum overlap length of 200 bp. Quality filters were used to remove sequences containing uncorrectable barcodes, ambiguous bases, or low quality reads (Phred quality scores ≤ 25). The open-reference OTU picking pipeline was performed at 97% identity using QIIME default parameters. The reference based OTU picking step and taxonomy assignment used the 97% OTU reference sequences from most recent release of the Greengenes database (August, 2013). In order to include as many samples and sequences as possible, the data was rarefied at 18,429 sequences per sample, which was the lowest sample sequencing depth over 1,000. The full dataset included 874 samples: 435 whole fecal microbiomes, 108 FACS Q1, 107 FACS Q2, 112 FACS Q3, and 112 FACS Q4.

Weighted and unweighted UniFrac beta diversity metrics [[4]](https://paperpile.com/c/IdncC4/aCRjd) were calculated using the rarefied OTUs. Principal coordinates analysis (PCoA) was performed on the weighted and unweighted UniFrac distance matrices of the full dataset as well as on subsets of the data for use in downstream analyses that examine only a portion of the samples. The reduced datasets include: (i) only the unsorted samples, (ii) only FACS samples, (iii) and the FACS samples excluding Q4. Alpha diversity metrics (Chao 1, observed number of OTUs, Faith’s phylogenetic diversity, and the Gini coefficient) for each sample were calculated as the average of estimates from 100 iterations of rarefaction at a depth of 18,429 sequences per sample.

**Statistical analyses**

*Defining the timepoint variable -* For some analyses and figures, a timepoint (e.g., T1, T2, T3, etc) category was necessary to allow for comparison of averages at each timepoint (see online supplementary figure S13). The timepoint assigned to a sample closely matches the age in months of the child at the time of sampling, however timepoint and age do not exactly match because of uneven times between samplings and dates across subjects. To assign a timepoint to a sample, first the age in months was calculated as the age in days divided by 30.5. Then the sample was assigned a timepoint by truncating the age in months at the time of sampling (e.g. 3.7 becomes T3). However, in the case where two consecutive samples were assigned the same timepoint (e.g., 3.1 and 3.9 are both assigned T3), the timepoint was adjusted so that the sample closest to the truncated month was assigned that timepoint, and the other sample was assigned the month it was closest to. As a result, no samples were assigned the same timepoint.

*Association of infant birth weight with mother BMI and gestational age -* A linear model with the response variable as the infant birth weight in grams (**BabyWeight**) was used to assess the effect of mother BMI (**MotherBMI**) and gestational age (**GestationalAge**) on the infant’s weight at birth. An interaction term for the interaction between mother BMI and gestational age (**MotherBMI**:**GestationalAge**) was also included in the model. The model below is written using notation from R.

**BabyWeight** ~ **MotherBMI** + **GestationalAge** + **MotherBMI**:**GestationalAge** + ε

*Association between stool IgA levels, age and breastfeeding status -* A linear mixed model was used to determine the impact of age (**Age**) and breastfeeding status (**Breastfeeding**) on IgA levels in the stool (**IgA**).

**IgA** ~ **Age** + **Breastfeeding** + **Age**:**Breastfeeding** + (1|**SUBJECT**) + ε

Age was represented in days and IgA levels were log transformed. A random effects term was used to account for the longitudinal sampling of subjects (1|**SUBJECT**).

*Association of subject characteristics with beta diversity and alpha diversity -* The linear mixed model below was used to determine which recorded participant characteristics are most significantly associated with the diversity of the infant fecal microbiomes. A model was fit for each of the top ten PCs from the PCoA of both unweighted UniFrac and weighted UniFrac and four alpha diversity metrics (Choa 1, Phylogenetic Diversity, Observed Species, and the Gini coefficient).

**y** ~ **Age + Geography** + **HasHadAntibiotic** + **HLA** + **Breastfeeding** + **DeliveryMode** + **BabyWeight** + **MotherBMI** + (1|**SUBJECT**) + ε

**y** is the PC or alpha diversity metric. The fixed effects in the model were age (**Age**), geographic location (**Geography**), a binary variable indicating if the infant has been given an oral antibiotic at any point prior to sampling (**HasHadAntibiotic**), HLA allele group (**HLA**), breastfeeding status (**Breastfeeding**), delivery mode (vaginal or cesarean; **DeliveryMode**), baby weight at delivery (**BabyWeight**), and mother’s BMI before pregnancy (**MotherBMI**). A random effects term was used to account for the longitudinal sampling of subjects (1|**SUBJECT**). The Benjamini-Hochberg procedure was used to correct for testing of the 20 PCs or the four alpha diversity metrics.

A similar procedure was applied to alpha and beta diversity in the FACS sorted microbiome with the following model specification.

**y** ~ **Age** + **Quadrant** + **Breastfeeding** + (1|**SUBJECT**) + (1|**SAMPLE**) + ε

The model included a fixed effect for each of the four (Q1-Q4) sorted populations (**Quadrant**) and random effect terms to account for the longitudinal sampling of subjects (1|**SUBJECT**) and the multiple unsorted or FACS populations coming from the same sample (1|**SAMPLE**). Due to the significantly lower number of cells found in Q4 across the samples, this same model was repeated with the exclusion of Q4.

*Association of stool levels of IgA, IgG, with common OTUs -* For each OTU shared by at least 40% of the samples, the association with IgA and IgG was assessed using the linear mixed model below which included age and breastfeeding status at the time of sampling. **y** ~ **Age** + **IgA** + **lgG** + **Breastfeeding** + **Age:Breastfeeding** + (1|**SUBJECT**) + ε

**y** is the log transformed rarefied OTU count with an offset of 1 applied. The fixed effects in the model were age in days (**Age**), log of stool IgA levels (**IgA**), log of stool IgG levels with an offset of 0.01 (**IgG**), breastfeeding status (**Breastfeeding**). A random effects term was used to account for the longitudinal sampling of subjects (1|**SUBJECT**). The Benjamini-Hochberg procedure was used to correct for testing of the 191 OTUs.

*Correlation between IgA, IgM, and IgG signals from FACS sorting of the fecal samples -* The package flowCore was used to extract the fluorescent intensities from the raw flow cytometry standard files. A pearson correlation was performed between the IgA and IgM signals and between IgA and IgG signals.

*Comparison of alpha diversity between unsorted microbiota and FACS quadrants -* A linear mixed model was fit for each of the four alpha diversity metrics (Choa 1, Phylogenetic Diversity, Observed Species, and the Gini coefficient).

**y** ~ **Age + Quadrant** + (1|**SUBJECT**) + (1|**SAMPLE**) + ε

**y** is the alpha diversity metric. The fixed effects in the model were age (**Age**) and quadrant where quadrant refers to unsorted and the four sorted populations (**Quadrant**). Random effects terms were used to account for the longitudinal sampling of subjects (1|**SUBJECT**) and the multiple unsorted or FACS populations coming from the same sample (1|**SAMPLE**).

*Identification of differentially abundant OTUs between FACS quadrants -* A model specification similar to that used for comparing alpha diversity between the quadrants was used to look for common OTUs (present in at least 40% of the FACS samples) that were differentially abundant between Q1-Q3 and Q2-Q3. The difference from the model above is that **y** is the log transformed rarefied OTU count with an offset of 1 applied and prior to running the model the data was subset to only the quadrants being compared.

**Supplementary Figure Legends**

**S1.** Percent relative abundance of the dominant bacterial phyla (A) and families (B) in the unsorted samples (top plot in each panel) and each of the four quadrants is plotted over time. Q1: IgA and IgM both high, IgG low; Q2: All high; Q3: All Low; Q4: IgG high, IgA and IgM both low.

**S2.** First two axes (PC1 and PC2) from principal coordinates analysis of the unweighted UniFrac distances between the unsorted fecal microbiome samples of 32 infants over the first couple years of life. Points are colored by participant characteristics and phylogenetic diversity. (A) Infant age in days at the time of sampling. (B) Geographic location of the TEDDY associated primary care center where samples were collected; this includes samples from Germany, Sweden, Georgia (USA) and Washington (USA). (C) Vaginal or cesarean delivery. (D) Baby birth weight in grams. (E) Body mass index of the mother before pregnancy. (F) HLA genotypes of the infant: *DR4-DQA1*030X-DQB1*0302 / DR3-DQA1*0501-DQB1*0201* (*DR3/4*), *DR4-DQA1*030X-DQB1*0302 / DR4-DQA1*030X-DQB1*0302* (*DR4/4*), and *DR3-DQA1*0501-DQB1*0201 / DR3-DQA1*0501-DQB1*0201* (*DR3/3*). (G) Breastfeeding status at the time of sampling. (H) Faith’s phylogenetic diversity. In all plots colored by a quantitative trait blue indicates lower values and red indicates higher values.

**S3.** First two axes (PC1 and PC2) from principal coordinates analysis of the weighted UniFrac distances between the unsorted fecal microbiome samples of 32 infants over the first couple years of life. Points are colored by participant characteristics and phylogenetic diversity. (A) Infant age in days at the time of sampling. (B) Geographic location of the TEDDY associated primary care center where samples were collected; this includes samples from Germany, Sweden, Georgia (USA) and Washington (USA). (C) Vaginal or cesarean delivery. (D) Baby birth weight in grams. (E) Body mass index of the mother before pregnancy. (F) HLA genotypes of the infant: *DR4-DQA1*030X-DQB1*0302 / DR3-DQA1*0501-DQB1*0201* (*DR3/4*), *DR4-DQA1*030X-DQB1*0302 / DR4-DQA1*030X-DQB1*0302* (*DR4/4*), and *DR3-DQA1*0501-DQB1*0201 / DR3-DQA1*0501-DQB1*0201* (*DR3/3*). (G) Breastfeeding status at the time of sampling. (H) Faith’s phylogenetic diversity. In all plots colored by a quantitative trait blue indicates lower values and red indicates higher values.

**S4.** (A-B) Principal coordinate 4 (PC4; A) or 7 (PC7; B) of unweighted UniFrac distances across timepoints. Lines show the mean PC values ± s.e.m. grouped by timepoint and geographic location (A) or previous antibiotic exposure (B). (C) Plot illustrating the time of stool collection and antibiotic use for each participant.

**S5.** Alpha diversity and the variance of alpha-diversity between subjects increases with age. Plots show Choa 1 (A), phylogenetic diversity (B), observed species (C), and Gini coefficient (D) over time. The blue line shows the mean diversity within each timepoint and grey shading shows two standard deviations from the timepoint mean. All plots only show unsorted 16S data.

**S6.** Levels of IgA, IgG, and IgM are all positively correlated with each other and negatively correlated with age and alpha diversity. The figure shows box-plots of the within-individual Pearson correlation coefficients between each of the following variables: Age, Unweighted UniFrac (PC1, PC2, and PC3), Phylogenetic diversity, IgA (log transformed), IgG (log transformed), and IgM (log transformed). The box-plots are colored by the repeated measures correlation (as calculated by rmcorr R package) and illustrate the agreement between the mean within-individual Pearson correlation coefficient and the repeated measures correlation for each pair of variables.

**S7.** Alpha diversity differs between the sorted and unsorted fraction of the gut microbiome. Each panel shows how the alpha-diversity metric (Choa 1: A, E; Phylogentic diversity: B, F; observed species: C, G; Gini coefficient: D, H) changes over time. The lines indicate the mean ± s.e.m. within a timepoint and either geographic location (A-D) or FACS quadrant (E-H).

**S8.** Different taxa are enriched in the sorted versus unsorted fractions. For each taxon the left plot is the relative abundance (log transformed) of the taxon over time, where the lines are the mean ± s.e.m. of the relative abundance within a timepoint and FACS quadrant (or the unsorted fecal microbiome). The plot on the right is a box-plot illustrating the overall difference in the relative abundance of the taxon (log transformed) between the sorted and unsorted fractions with all timepoints combined. The title of each plot is the taxon and the nominal p value of the enrichment in the sorted compared to the unsorted fractions (see methods) and the plots are arranged by their significance, with more significant taxa at the top left of the figure.

**S9.** Beta diversity analysis of the sorted fecal microbiome. First two axes (PC1 and PC2) from principal coordinates analysis of the unweighted (panels A-E) and weighted (panels F-J) UniFrac distances between the sorted fecal microbiome samples of 32 infants over the first couple years of life. Points are colored by participant characteristics and phylogenetic diversity. (A, F) Anonymous sample ID. (B,G) FACS quadrant. (C,H) Infant age in days at sample collection. (D, I) Faith’s phylogenetic diversity. (E, J) Whether the infant was currently receiving breastmilk.

**S10.** The percentage of the sorted microbiome fraction by FACS quadrant. For each subject we quantified the percentage of the sorted fecal microbiome that was detected in each of the 4 quadrants. Subjects are listed along the y-axis, with relative abundance shown along the x-axis, and colored by FACS quadrant.

**S11.** All differentially abundant OTUs between Q1 (IgM and IgA both high) and Q3 (low coating)*.* Plots on the left show the log(Q1 OTU abundance) - log(Q3 OTU abundance) divided by log(Q1 OTU abundance) + log(Q3 OTU abundance) over time where each point represents a sample and is colored by timepoint. Positive values on the y-axis indicate enrichment of the OTU abundance in Q1 and negative values indicate and enrichment of the OTU abundance in Q3; this is similar to the IgA index defined in Planer et al 2016. Plots on the right show the average of the log transformed OTU relative abundance in Q1 (Red) and Q3 (Blue) at each time point. OTU Greengenes ID, taxonomic classification, and the p value from the linear mixed model using OTU as a response variable is indicated above each set of graphs. Only common OTUs (non zero value in >40% of samples tested) were tested. The plots are arranged by their significance, with more significant OTUs at the top left of the figure.

**S12.** Same plots as S10 but showing differentially expressed OTUs between Q2 (all high coating) and Q3 (all low coating).

**S13.** Illustration of the relationship between the timepoint variable and the age of the participants. Sample collection from each participant started around 2-3 months of age and subsequent samples were collected approximately each month following the initial sample. Some analyses relied on defining timepoints that relate to the age of the participant at the time of sampling and are also comparable across participants. The full method for assigning timepoints to a sample is described in the methods. (A-B) In both plots, each point indicates a stool sample and the x-axis shows the participants age in days at the time of sample collection. (A) The y-axis is the timepoint variable used throughout the analyses, we see that there is minimal overlap in age between the timepoints. (B) The y-axis is the participant's anonymized ID, and this plot shows that across participants each timepoint reflects a similar age.

**The TEDDY Study Group**

**Colorado Clinical Center:** Marian Rewers, M.D., Ph.D., PI^1,4,5,6,9,10^, Aaron Barbour, Kimberly Bautista^11^, Judith Baxter^8,911^, Daniel Felipe-Morales, Kimberly Driscoll, Ph.D.^8^, Brigitte I. Frohnert, M.D.^2,13^, Marisa Stahl, M.D.^12^, Patricia Gesualdo^2,6,11,13^, Michelle Hoffman^11,12,13^, Rachel Karban^11^, Edwin Liu, M.D.^12^, Jill Norris, Ph.D.^2,3,11^, Stesha Peacock, Hanan Shorrosh, Andrea Steck, M.D.^3,13^, Megan Stern, Erica Villegas^2^, Kathleen Waugh^6,7,11^. University of Colorado, Anschutz Medical Campus, Barbara Davis Center for Childhood Diabetes.

**Finland Clinical Center:** Jorma Toppari, M.D., Ph.D., PI^¥^1,4,10,13^, Olli G. Simell, M.D., Ph.D., Annika Adamsson, Ph.D.^^11^, Suvi Ahonen*^±§^, Mari Åkerlund*^±§^, Leena Hakola*, Anne Hekkala, M.D.^µ¤^, Henna Holappa^µ¤^, Heikki Hyöty, M.D., Ph.D.*^±6^, Anni Ikonen^µ¤^, Jorma Ilonen, M.D., Ph.D.^¥¶3^, Sinikka Jäminki*^±^, Sanna Jokipuu^^^, Leena Karlsson^^^, Jukka Kero M.D., Ph.D.^¥^^, Miia Kähönen^µ¤11,13^, Mikael Knip, M.D., Ph.D.*^±5^, Minna-Liisa Koivikko^µ¤^, Merja Koskinen*^±^, Mirva Koreasalo*^±§2^, Kalle Kurppa, M.D., Ph.D.*^±12^, Jarita Kytölä*^±^, Tiina Latva-aho^µ¤^, Katri Lindfors, Ph.D.*^12^, Maria Lönnrot, M.D., Ph.D.*^±6^, Elina Mäntymäki^^^, Markus Mattila*, Maija Miettinen^§2^, Katja Multasuo^µ¤^, Teija Mykkänen^µ¤^, Tiina Niininen^±^*^11^, Sari Niinistö^±§2^, Mia Nyblom*^±^, Sami Oikarinen, Ph.D.*^±^, Paula Ollikainen^µ¤^ , Zhian Othmani^^^, Sirpa Pohjola ^µ¤^, Petra Rajala^^^, Jenna Rautanen^±§^, Anne Riikonen*^±§2^, Eija Riski^^^, Miia Pekkola^*±^, Minna Romo^^^, Satu Ruohonen^^^, Satu Simell, M.D., Ph.D.^¥12^, Maija Sjöberg^^^, Aino Stenius^µ¤11^, Päivi Tossavainen, M.D.^µ¤^, Mari Vähä-Mäkilä^¥^, Sini Vainionpää^^^, Eeva Varjonen^^11^, Riitta Veijola, M.D., Ph.D.^µ¤13^, Irene Viinikangas^µ¤^, Suvi M. Virtanen, M.D., Ph.D.*^±§2^. ^¥^University of Turku, *Tampere University, ^µ^University of Oulu, ^^^Turku University Hospital, Hospital District of Southwest Finland, ^±^Tampere University Hospital, ^¤^Oulu University Hospital, §National Institute for Health and Welfare, Finland, ^¶^University of Kuopio.

**Georgia/Florida Clinical Center:** Jin-Xiong She, Ph.D., PI^1,3,4,10^, Desmond Schatz, M.D.*^4,5,7,8^, Diane Hopkins^11^, Leigh Steed^11,12,13^, Jennifer Bryant^11^, Katherine Silvis^2^, Michael Haller, M.D.*^13^, Melissa Gardiner^11^, Richard McIndoe, Ph.D., Ashok Sharma, Stephen W. Anderson, M.D.^^^, Laura Jacobsen, M.D.*^13^, John Marks, DHSc.*^11,13^, P.D. Towe*. Center for Biotechnology and Genomic Medicine, Augusta University. *University of Florida, ^^^Pediatric Endocrine Associates, Atlanta.

**Germany Clinical Center:** Anette G. Ziegler, M.D., PI^1,3,4,10^, Ezio Bonifacio Ph.D.*^5^, Anita Gavrisan, Cigdem Gezginci, Anja Heublein, Verena Hoffmann, Ph.D.^2^, Sandra Hummel, Ph.D.^2^, Andrea Keimer^¥2^, Annette Knopff^7^, Charlotte Koch, Sibylle Koletzko, M.D.^¶12^, Claudia Ramminger^11^, Roswith Roth, Ph.D.^8^, Marlon Scholz, Joanna Stock^8,11,13^, Katharina Warncke, M.D.^13^, Lorena Wendel, Christiane Winkler, Ph.D.^2,11^. Forschergruppe Diabetes e.V. and Institute of Diabetes Research, Helmholtz Zentrum München, Forschergruppe Diabetes, and Klinikum rechts der Isar, Technische Universität München. *Center for Regenerative Therapies, TU Dresden, ^¶^Dr. von Hauner Children’s Hospital, Department of Gastroenterology, Ludwig Maximillians University Munich, ^¥^University of Bonn, Department of Nutritional Epidemiology.

**Sweden Clinical Center:** Åke Lernmark, Ph.D., PI^1,3,4,5,6,8,9,10^, Daniel Agardh, M.D., Ph.D.^6,12^, Carin Andrén Aronsson, Ph.D.^2,11,12^, Maria Ask, Rasmus Bennet, Corrado Cilio, Ph.D., M.D.^5,6^, Helene Engqvist, Emelie Ericson-Hallström, Annika Fors, Lina Fransson, Thomas Gard, Monika Hansen, Hanna Jisser, Fredrik Johansen, Berglind Jonsdottir, M.D., Ph.D.^11^, Silvija Jovic, Helena Elding Larsson, M.D., Ph.D.^6,13^, Marielle Lindström, Markus Lundgren, M.D., Ph.D.^13^, Marlena Maziarz, Ph.D., Maria Månsson-Martinez, Maria Markan, Jessica Melin^11^, Zeliha Mestan, Caroline Nilsson, Karin Ottosson, Kobra Rahmati, Anita Ramelius, Falastin Salami, Anette Sjöberg, Birgitta Sjöberg, Malin Svensson, Carina Törn, Ph.D.^3^, Anne Wallin, Åsa Wimar^13^, Sofie Åberg. Lund University.

**Washington Clinical Center:** William A. Hagopian, M.D., Ph.D., PI^1,3,4,5,6,7,10,12,13^, Michael Killian^6,7,11,12^, Claire Cowen Crouch^11,13^, Jennifer Skidmore^2^, Masumeh Chavoshi, Rachel Hervey, Rachel Lyons, Arlene Meyer, Denise Mulenga^11^, Jared Radtke, Matei Romancik, Davey Schmitt, Sarah Zink. Pacific Northwest Research Institute.

**Pennsylvania Satellite Center:** Dorothy Becker, M.D., Margaret Franciscus, MaryEllen Dalmagro-Elias Smith^2^, Ashi Daftary, M.D., Mary Beth Klein, Chrystal Yates. Children’s Hospital of Pittsburgh of UPMC.

**Data Coordinating Center:** Jeffrey P. Krischer, Ph.D.,PI^1,4,5,9,10^, Sarah Austin-Gonzalez, Maryouri Avendano, Sandra Baethke, Rasheedah Brown^11^, Brant Burkhardt, Ph.D.^5,6^, Martha Butterworth^2^, Joanna Clasen, David Cuthbertson, Stephen Dankyi, Christopher Eberhard, Steven Fiske^8^, Jennifer Garmeson, Veena Gowda, Kathleen Heyman, Belinda Hsiao, Christina Karges, Francisco Perez Laras, Hye-Seung Lee, Ph.D.^1,2,3,12^, Qian Li^2,3^, Shu Liu, Xiang Liu, Ph.D.^2,3813^, Kristian Lynch, Ph.D. ^5,6,8^, Colleen Maguire, Jamie Malloy, Cristina McCarthy^11^, Aubrie Merrell, Hemang Parikh, Ph.D.^3^, Ryan Quigley, Cassandra Remedios, Chris Shaffer, Laura Smith, Ph.D.^8,11^, Susan Smith^11^, Noah Sulman, Ph.D., Roy Tamura, Ph.D.^1,2,11,12,13^, Dena Tewey, Michael Toth, Ulla Uusitalo, Ph.D.^2^, Kendra Vehik, Ph.D.^4,5,6,8,13^, Ponni Vijayakandipan, Keith Wood, Jimin Yang, Ph.D., R.D.^2^. *Past staff: Michael Abbondondolo, Lori Ballard, David Hadley, Ph.D., Wendy McLeod, Steven Meulemans.* University of South Florida.

**Project scientist:** Beena Akolkar, Ph.D.^1,3,4,5,6,7,9,10^. National Institutes of Diabetes and Digestive and Kidney Diseases.

**Other contributors:** Kasia Bourcier, Ph.D.^5^, National Institutes of Allergy and Infectious Diseases. Thomas Briese, Ph.D.^6^, Columbia University. Suzanne Bennett Johnson, Ph.D.^8,11^, Florida State University. Eric Triplett, Ph.D.^6^, University of Florida.

***Committees:***

^1^Ancillary Studies, ^2^Diet, ^3^Genetics, ^4^Human Subjects/Publicity/Publications, ^5^Immune Markers, ^6^Infectious Agents, ^7^Laboratory Implementation, ^8^Psychosocial, ^9^Quality Assurance, ^10^Steering, ^11^Study Coordinators, ^12^Celiac Disease, ^13^Clinical Implementation.
