## Supplementary figures and images for "Interactions between the gut microbiome and mucosal immunoglobulins A, M and G in the developing infant gut"

### Figure S1

**Supplementary figure S1**

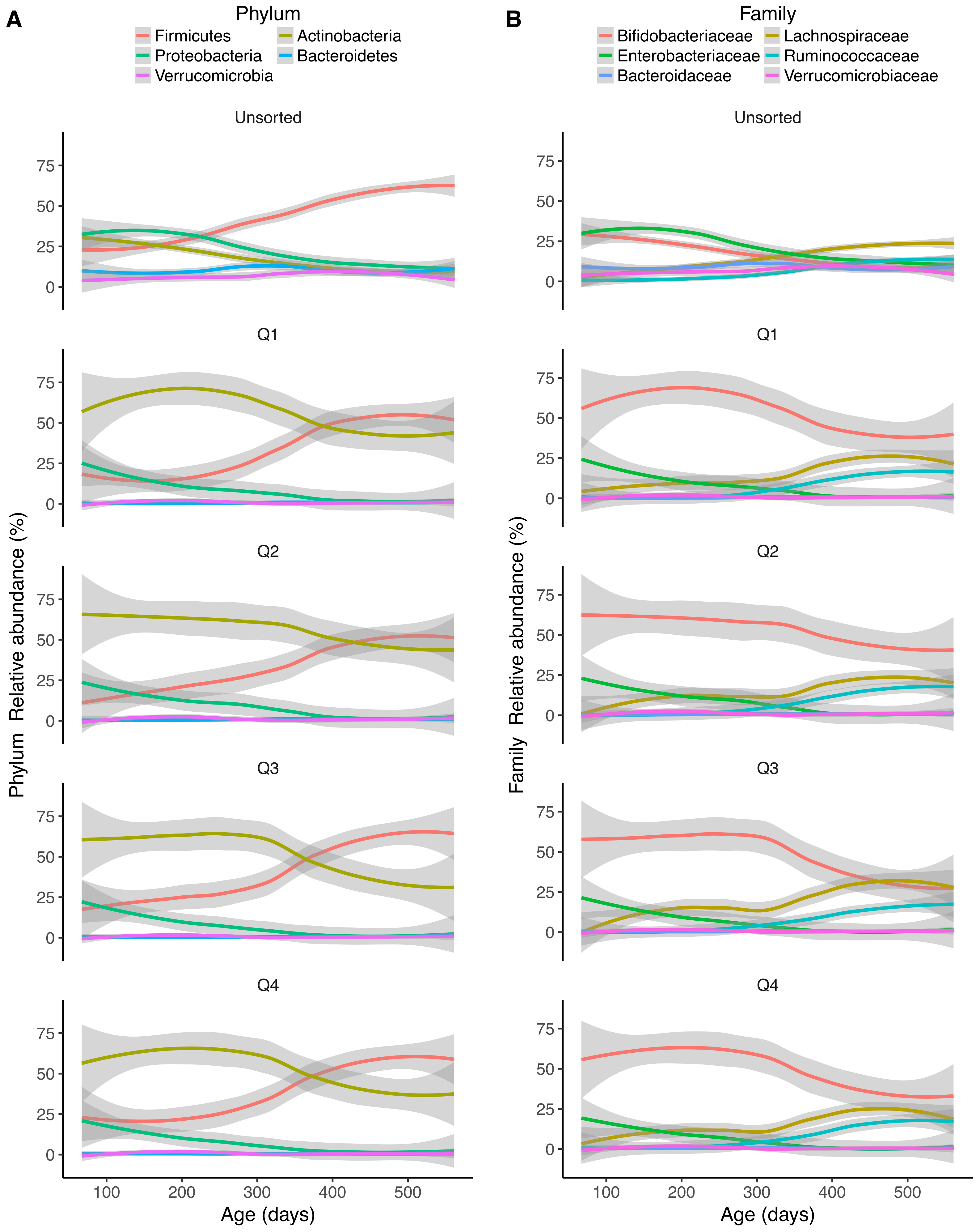

### Figure S2

#### Supplementary figure S2

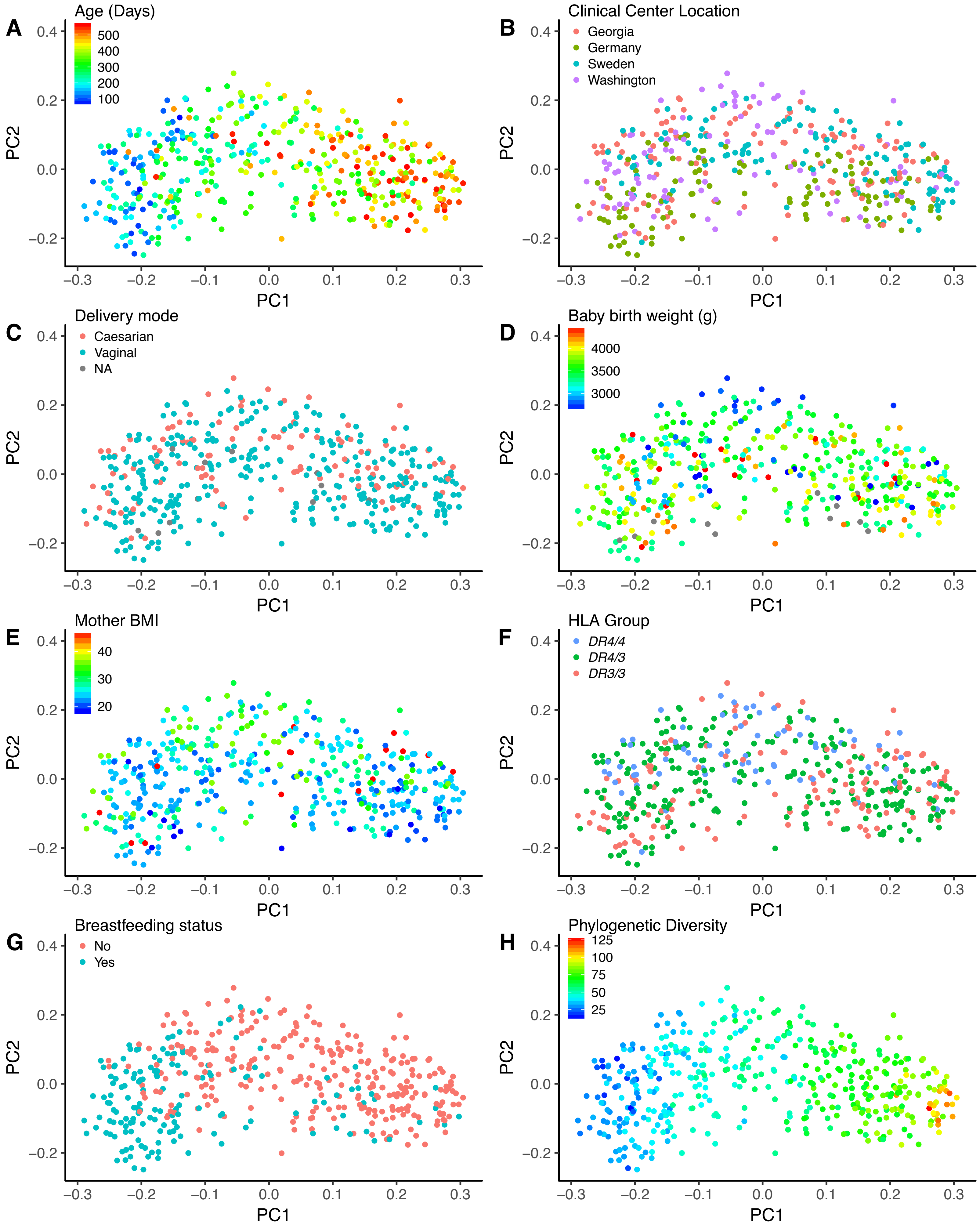

### Figure S3

##### Supplementary figure S3

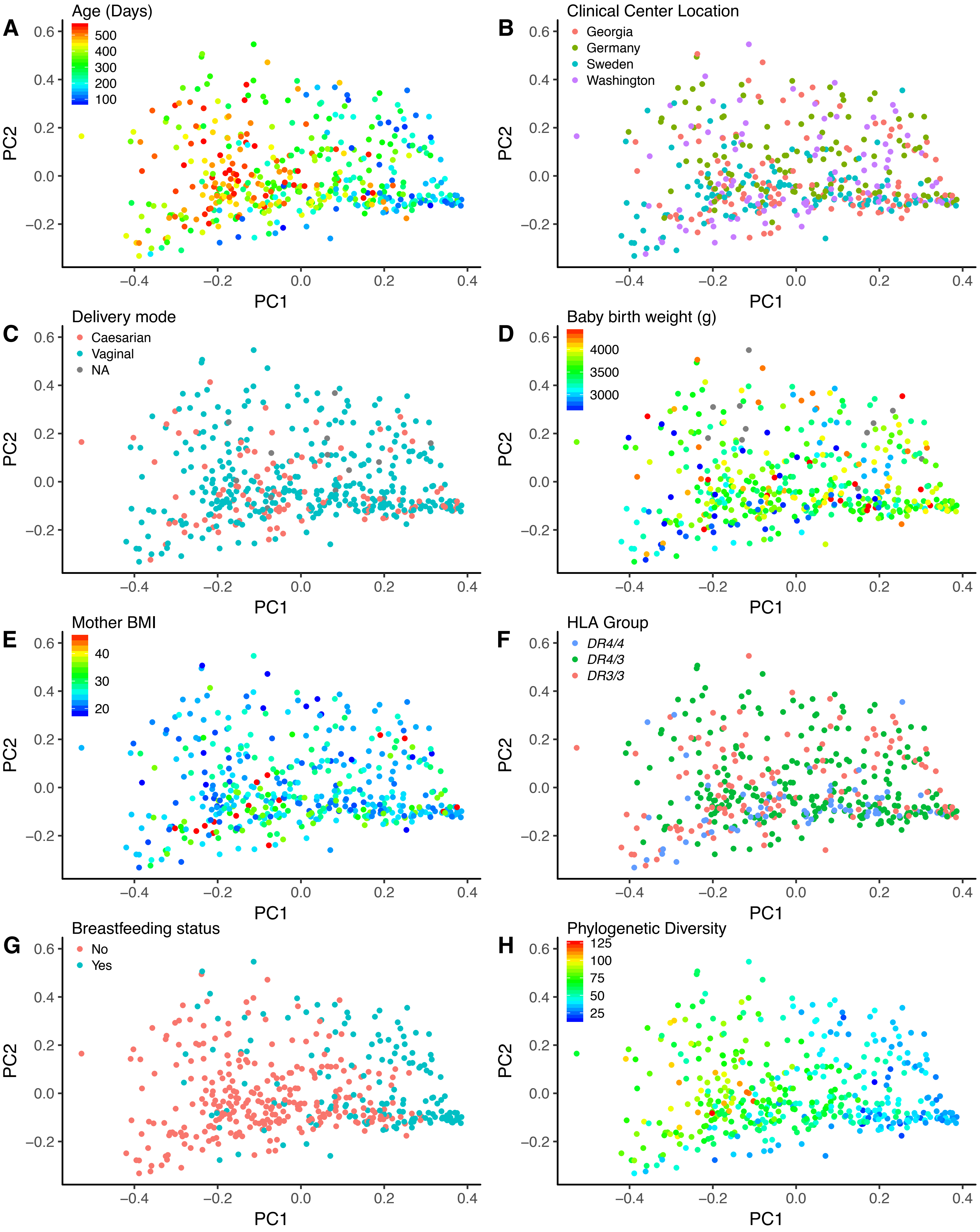

### Figure S4

##### Supplementary figure S4

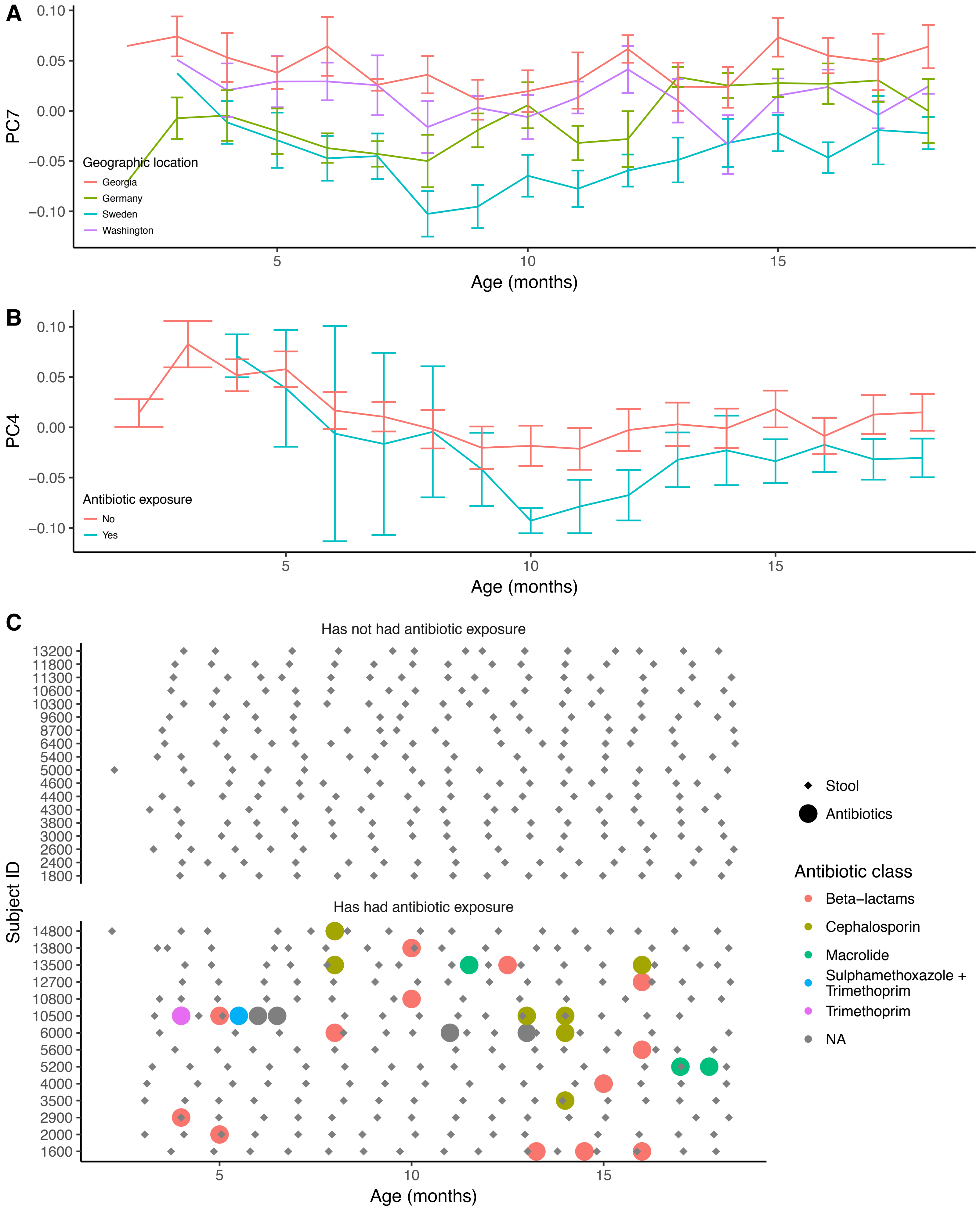

### Figure S5

#### Supplementary figure S5

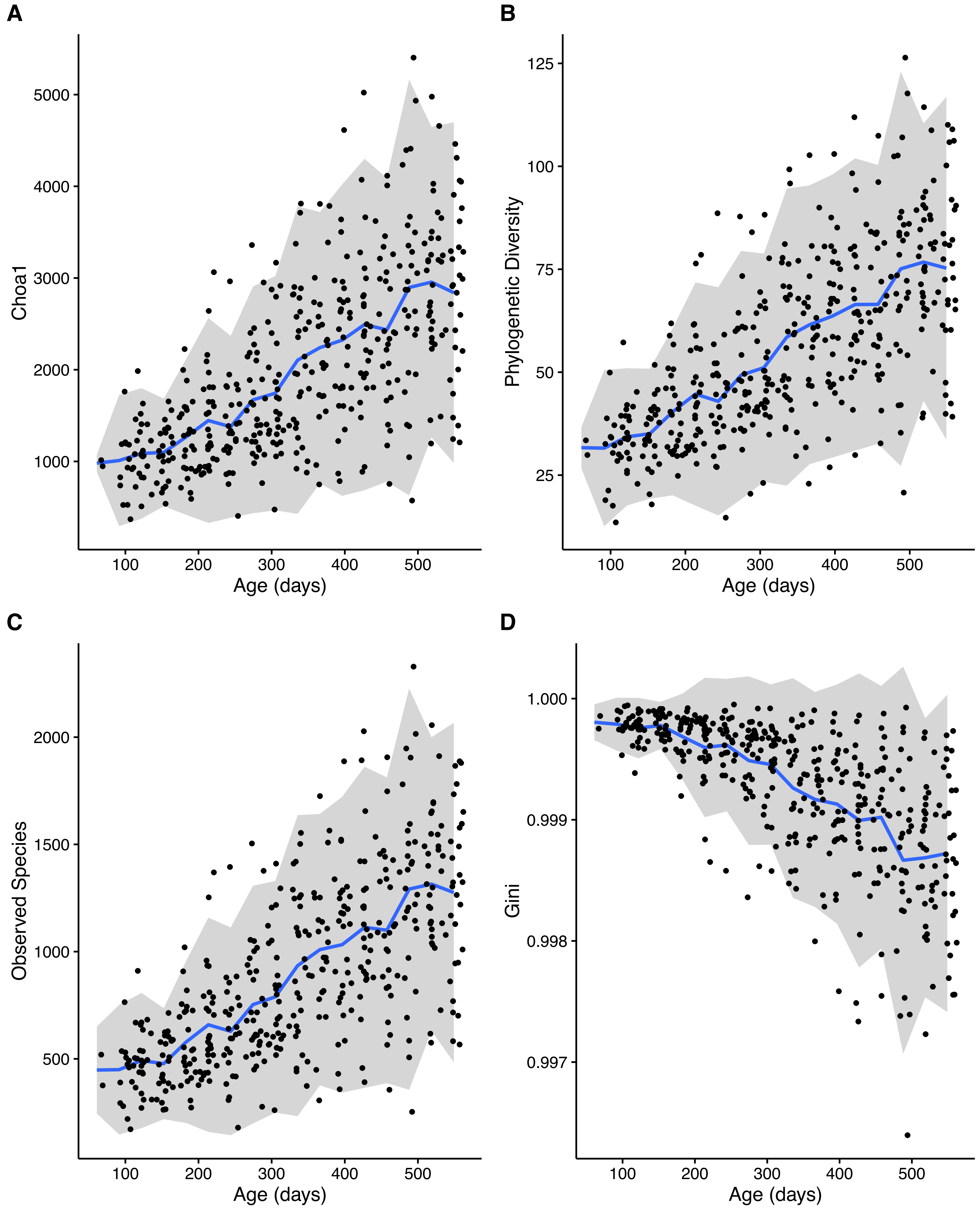

### Figure S6

**Supplementary figure S6**

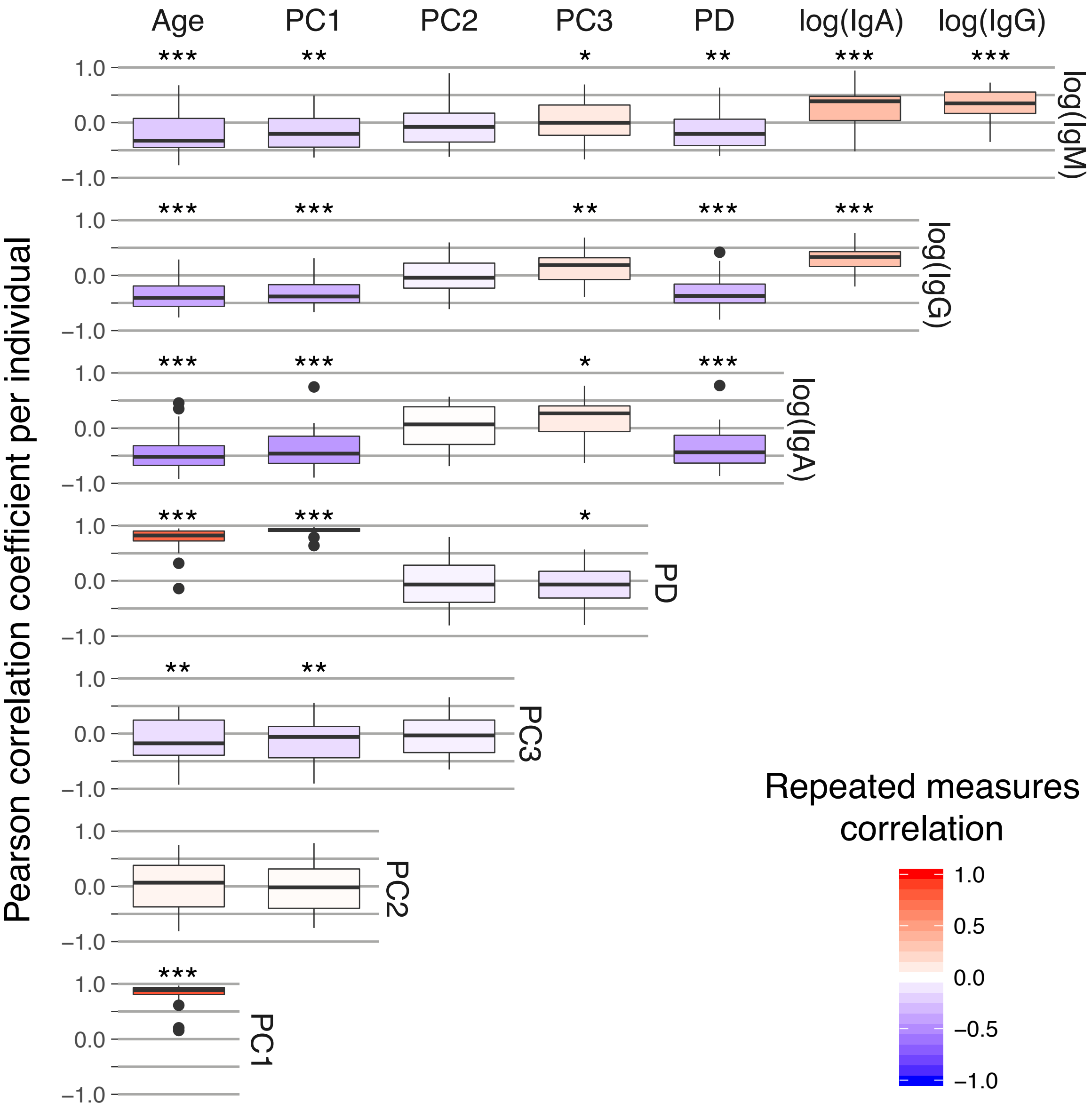

### Figure S7

#### Supplementary figure S7

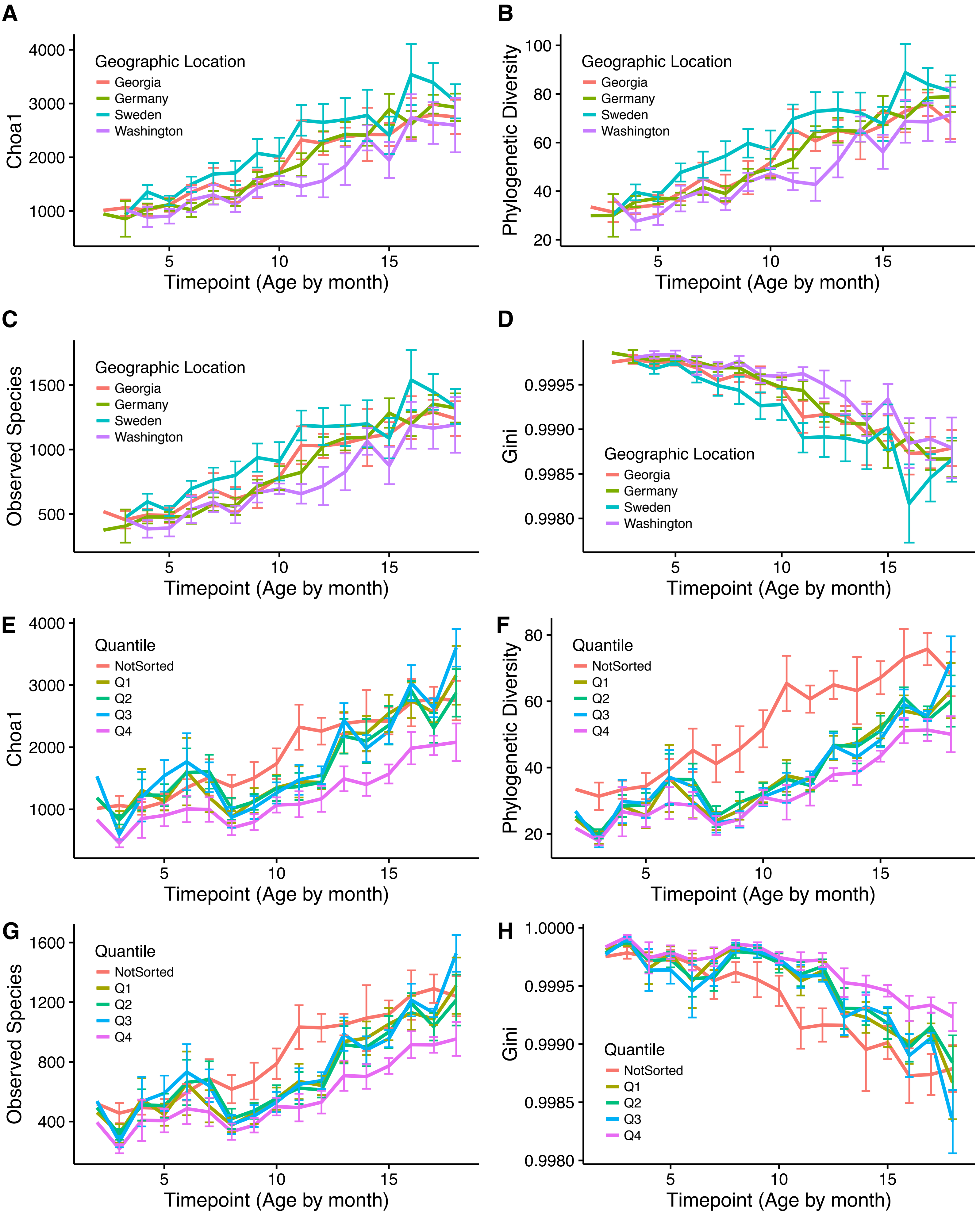

### Figure S8

#### Supplementary figure S8

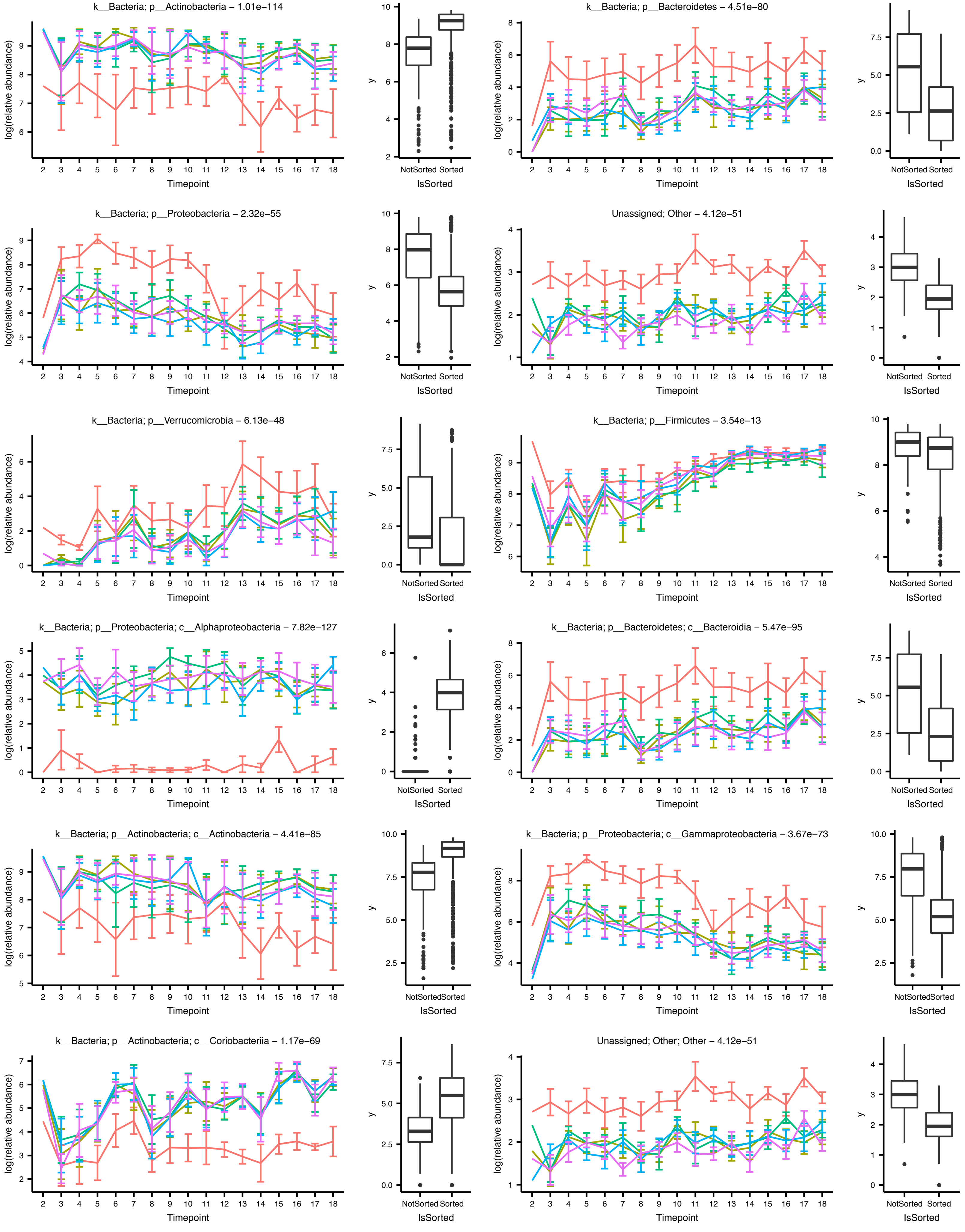

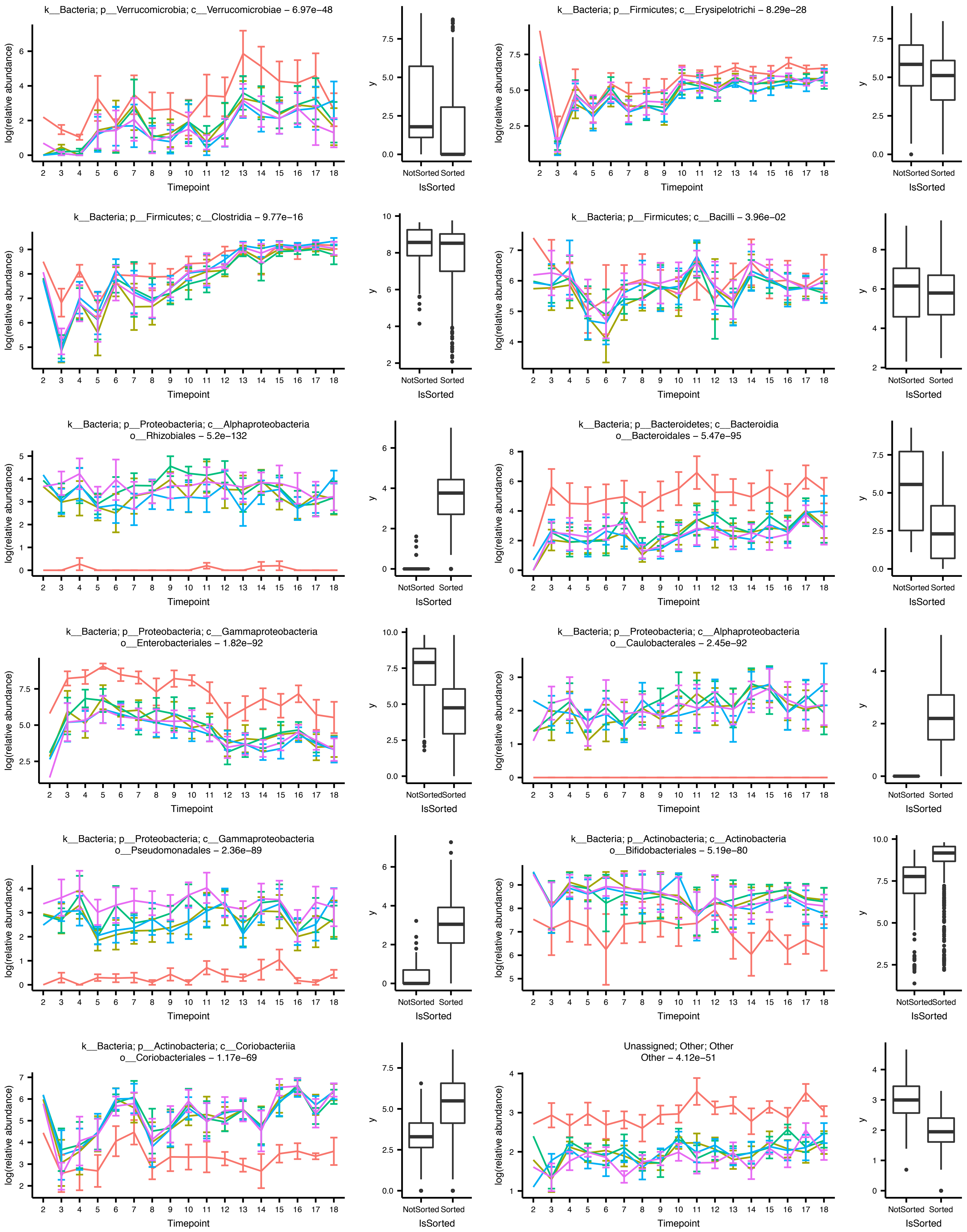

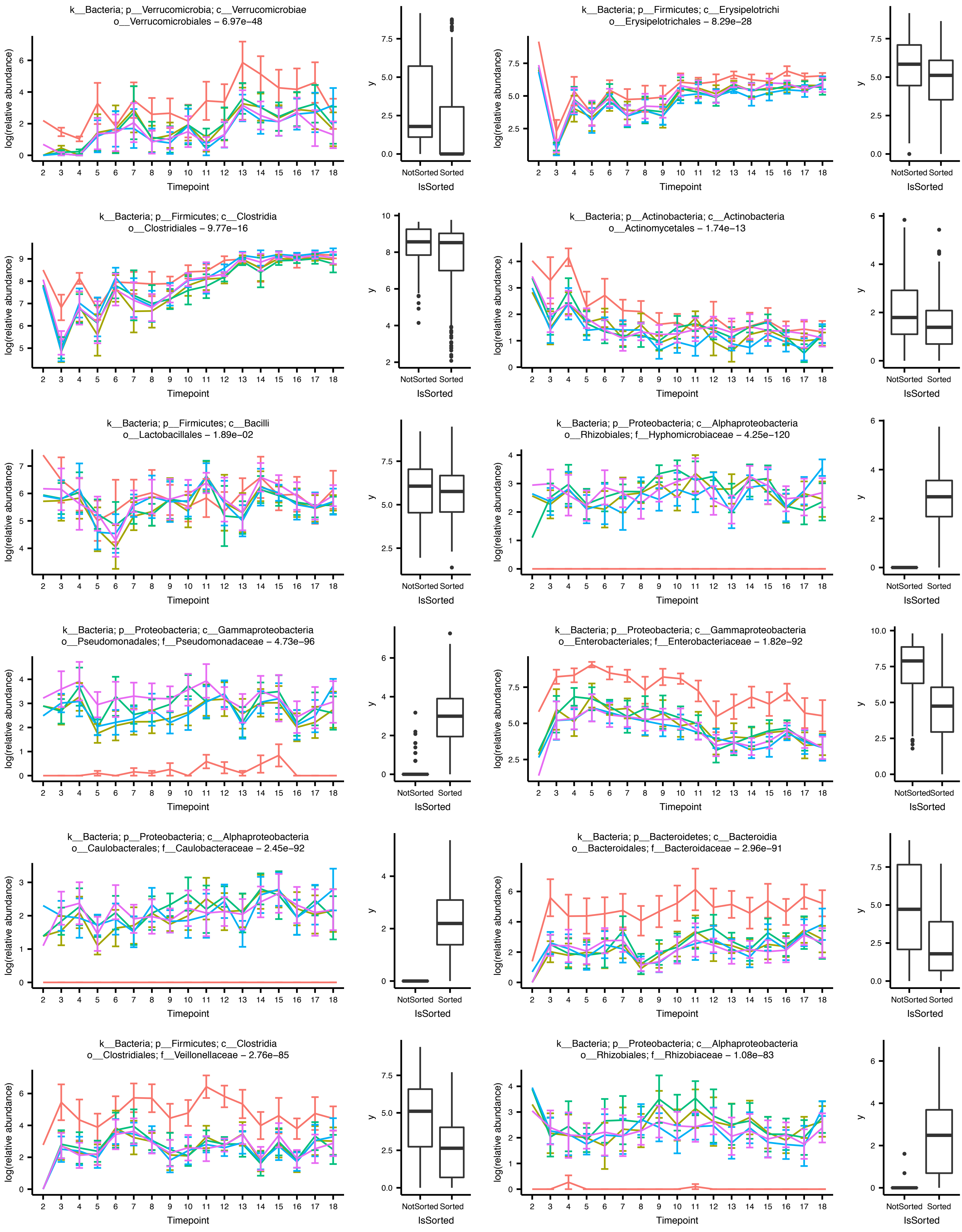

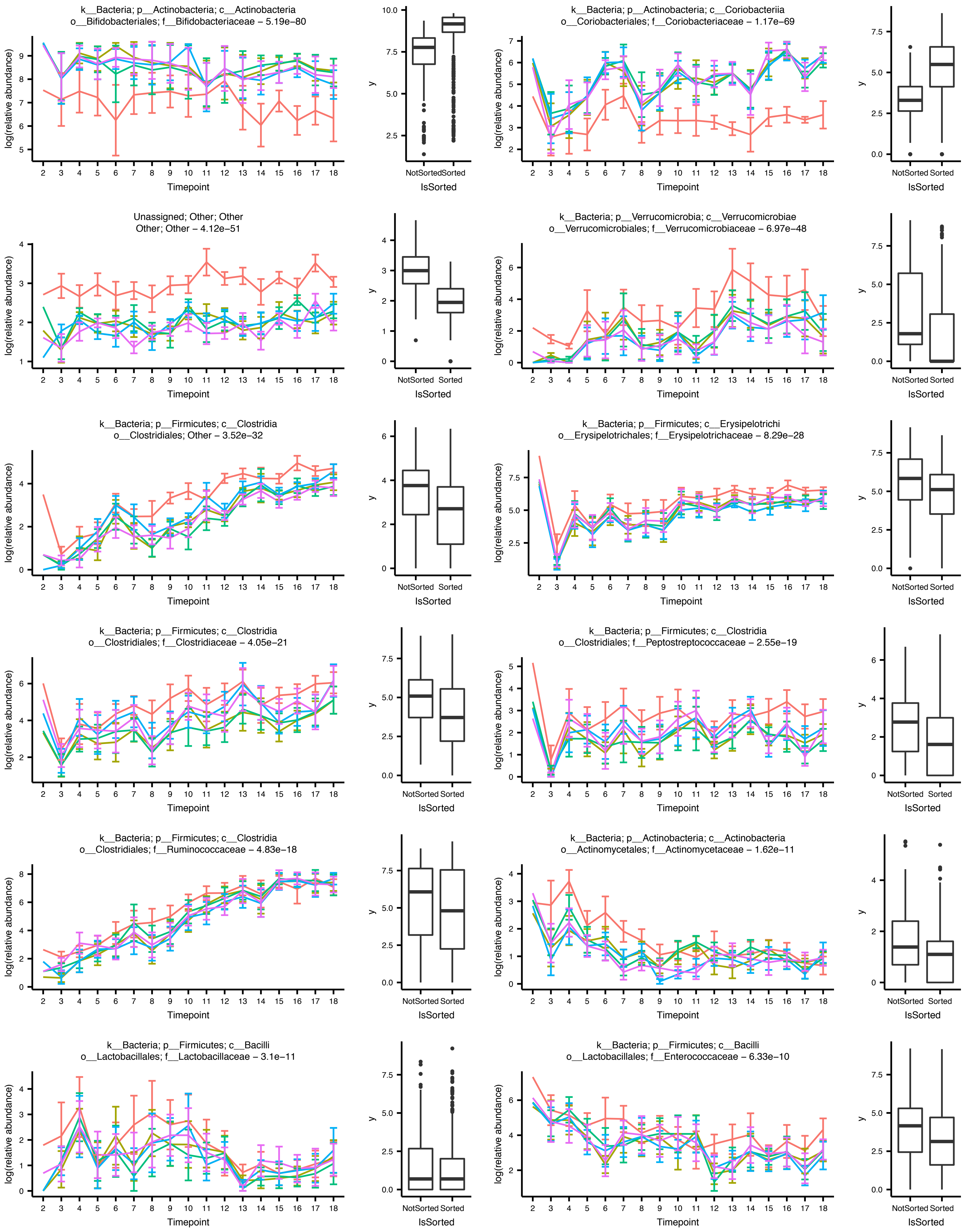

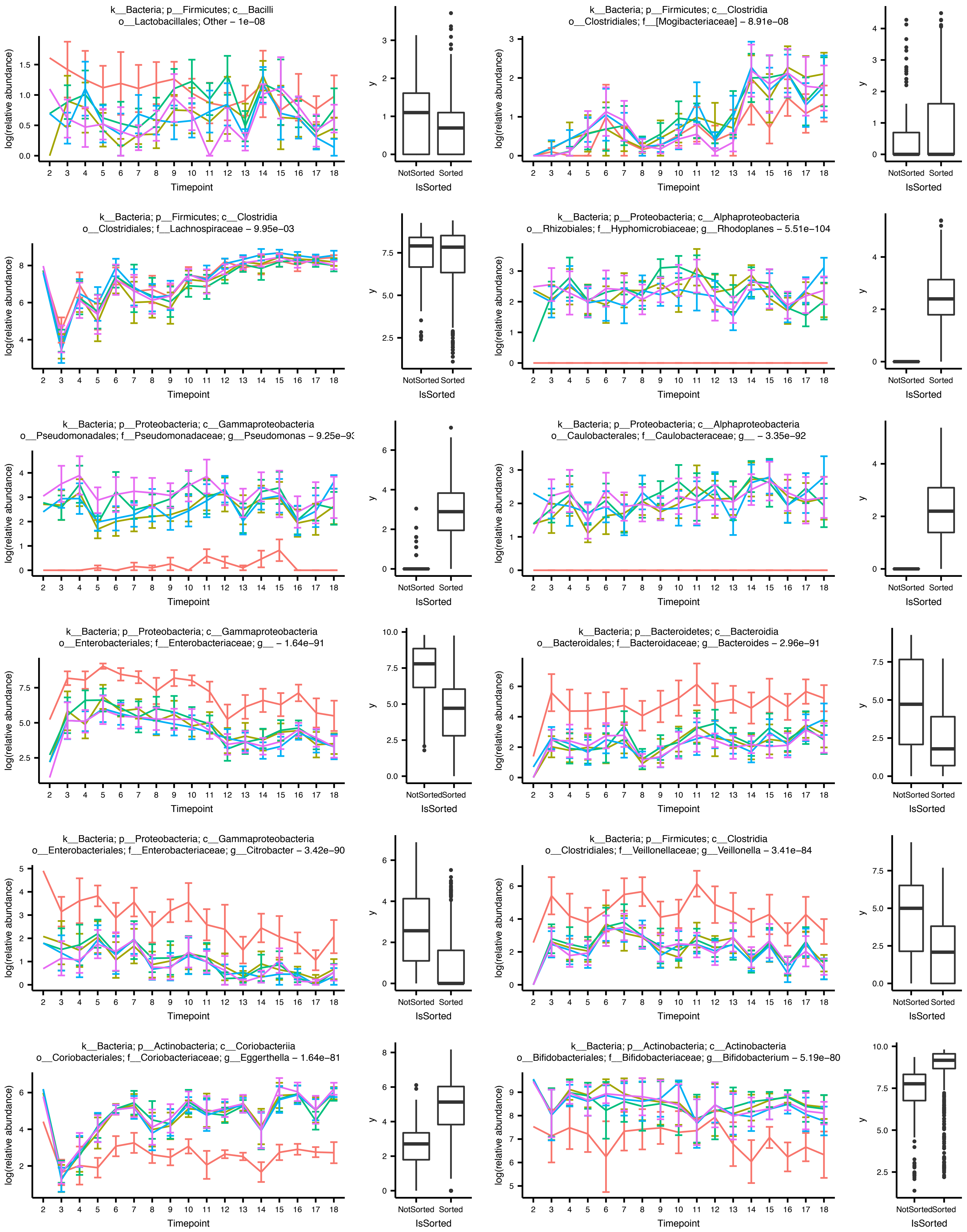

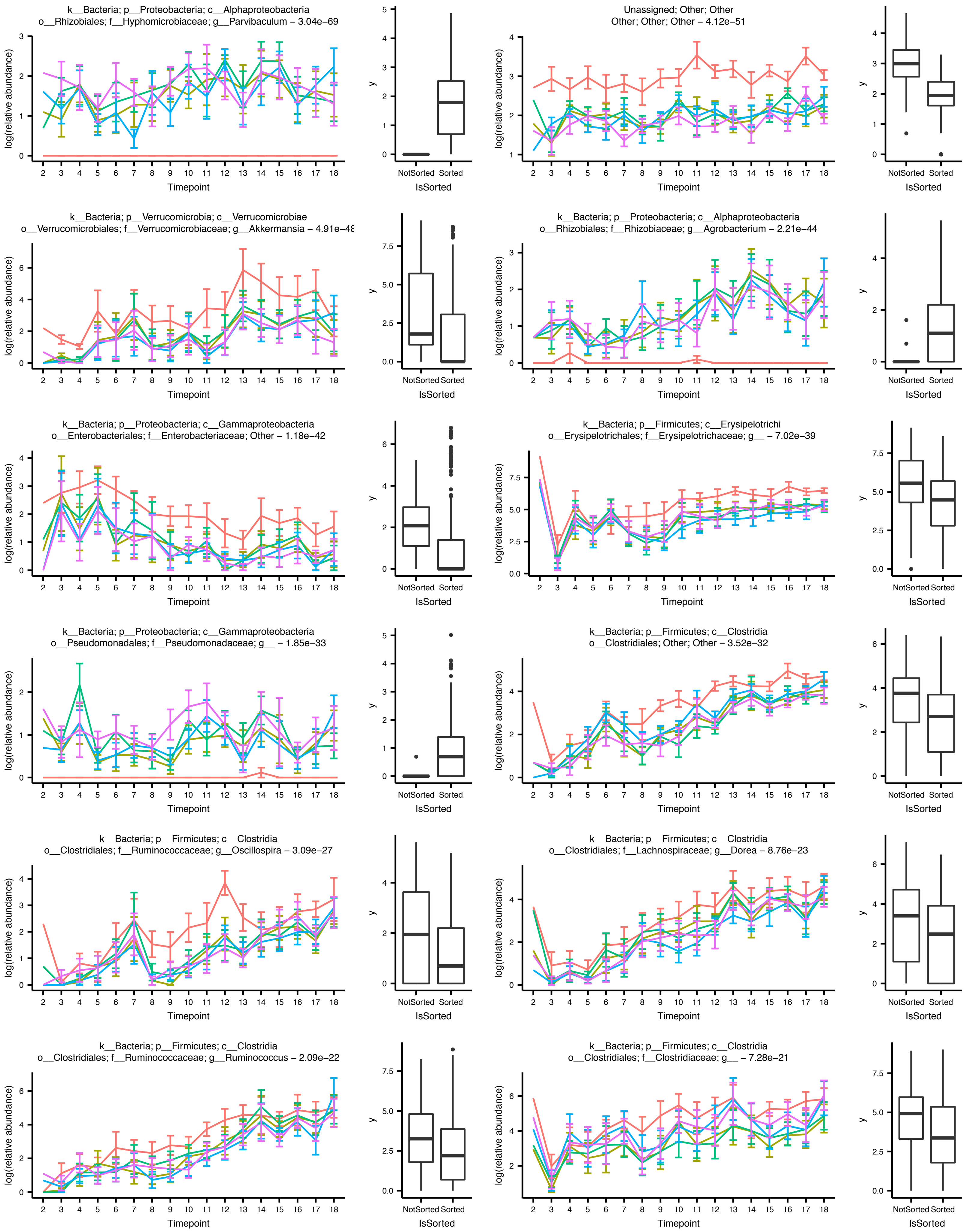

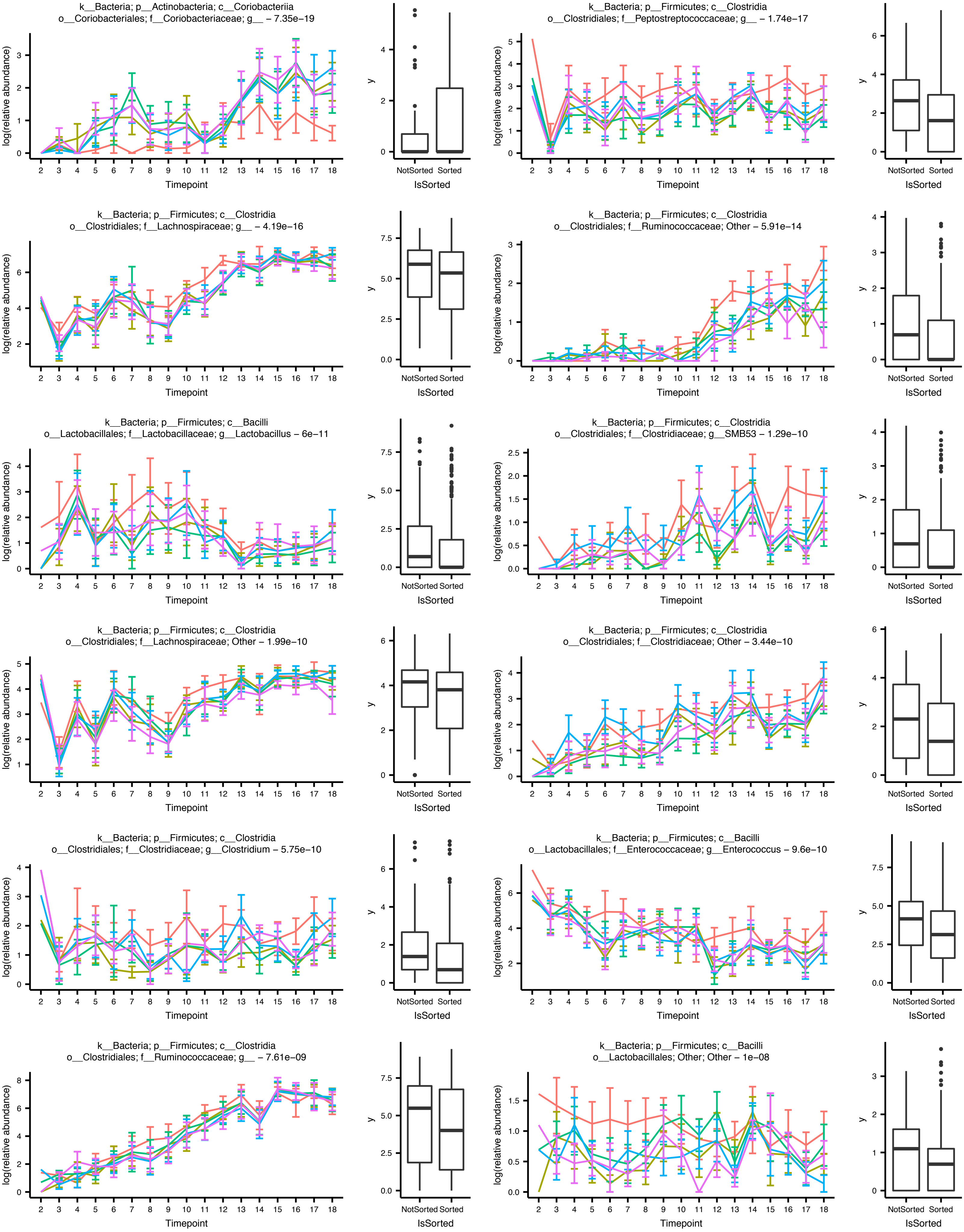

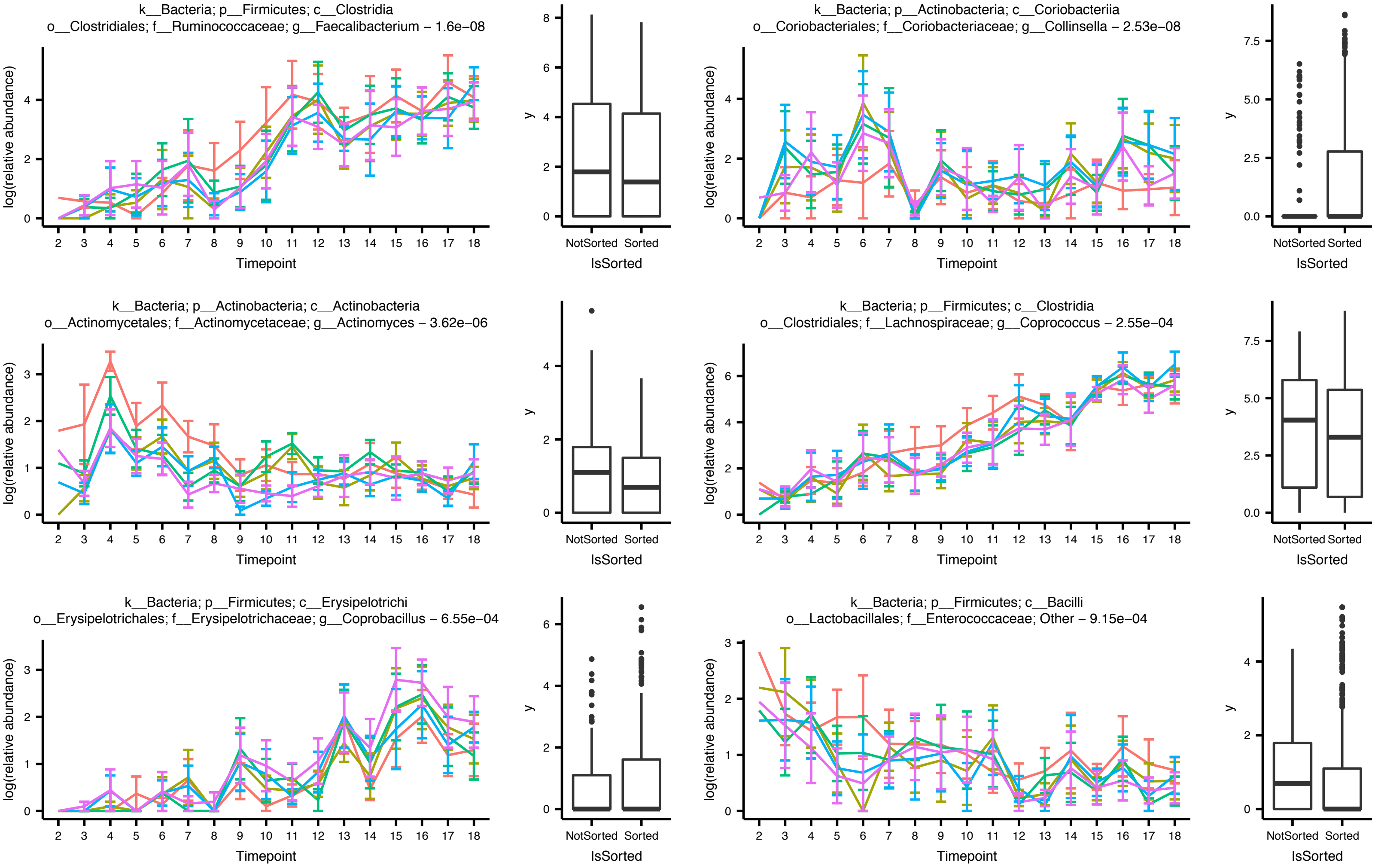

### Figure S9

#### Supplementary figure S9

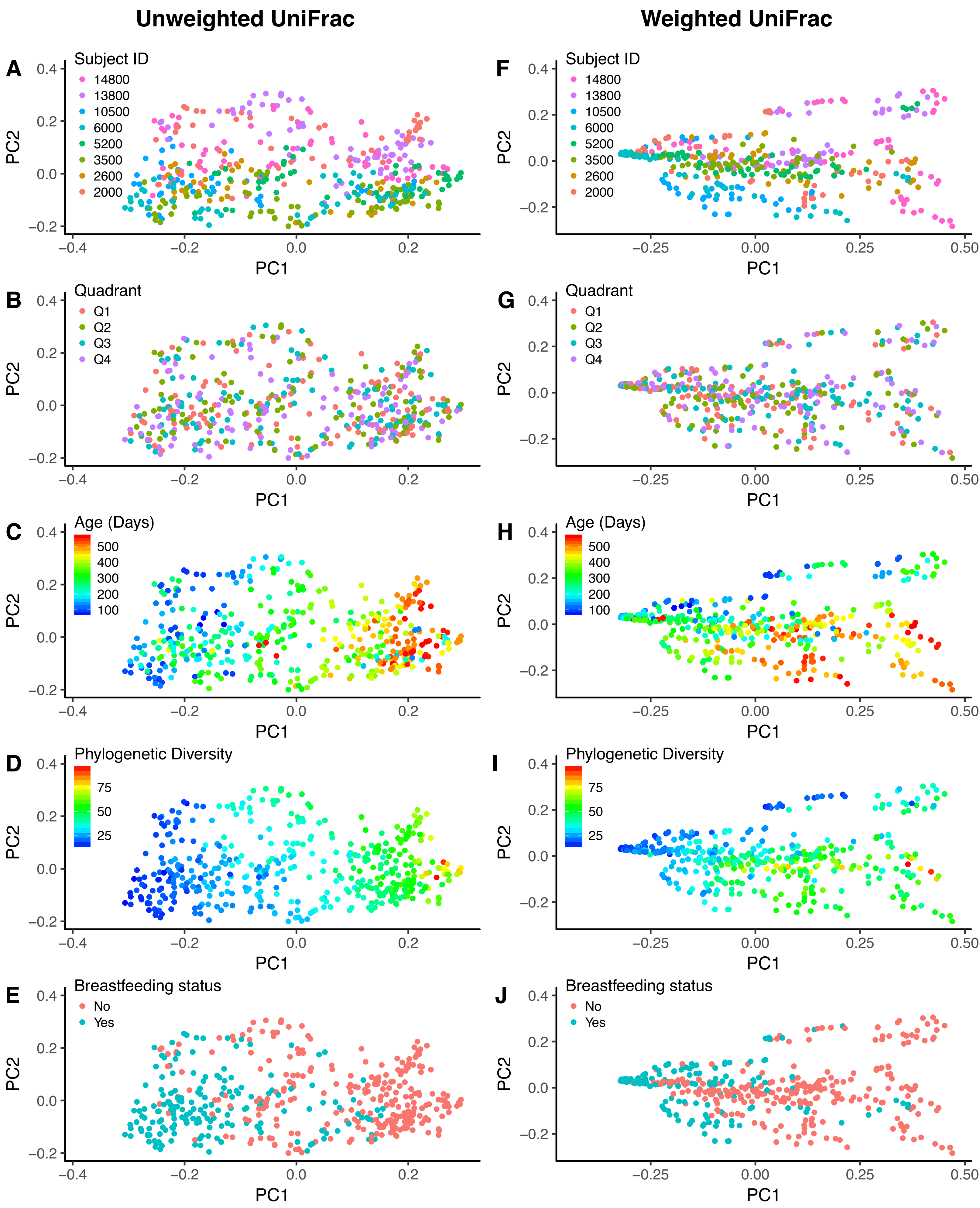

### Figure S10

#### Supplementary figure S10

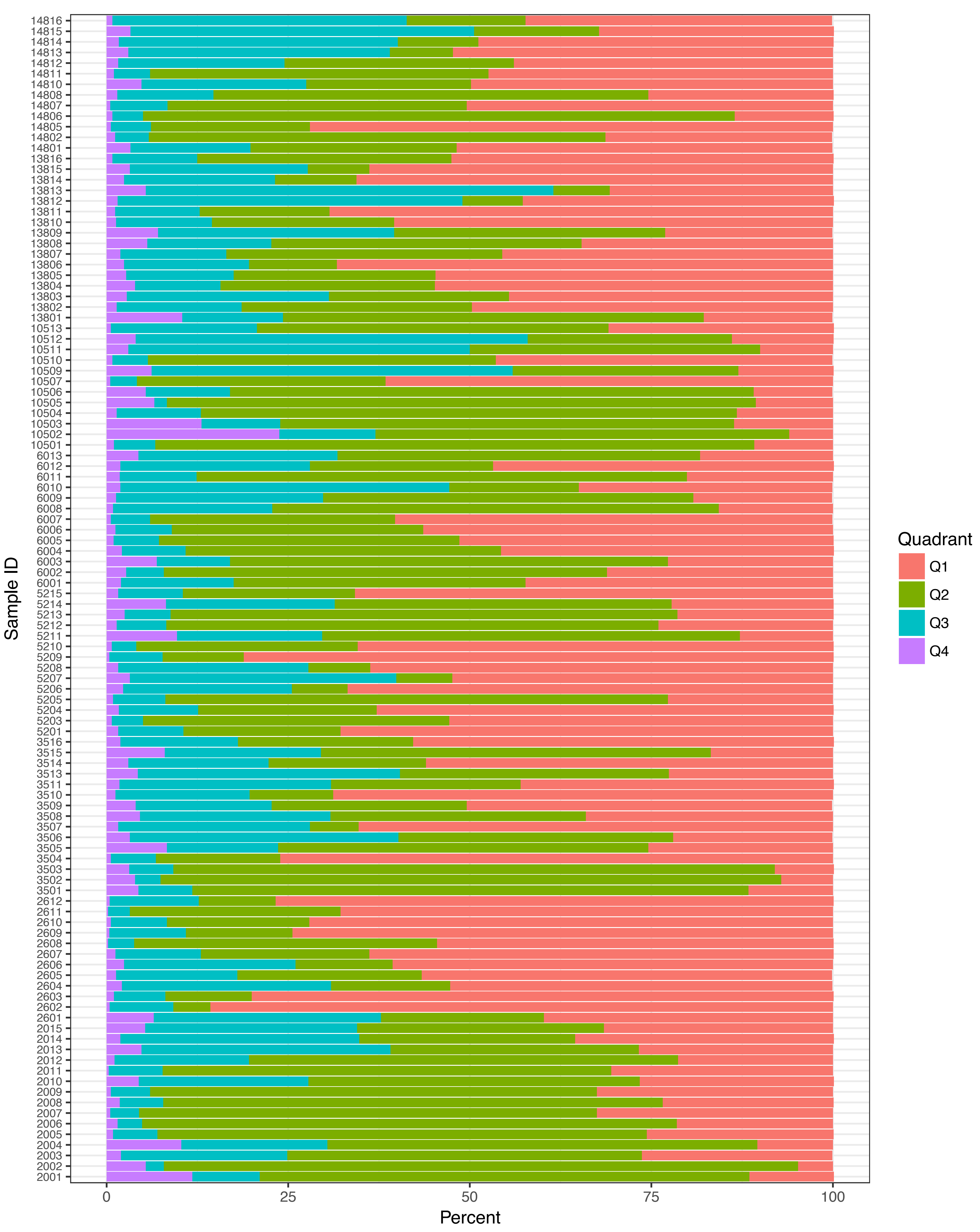

### Figure S13

**Supplementary figure S13**

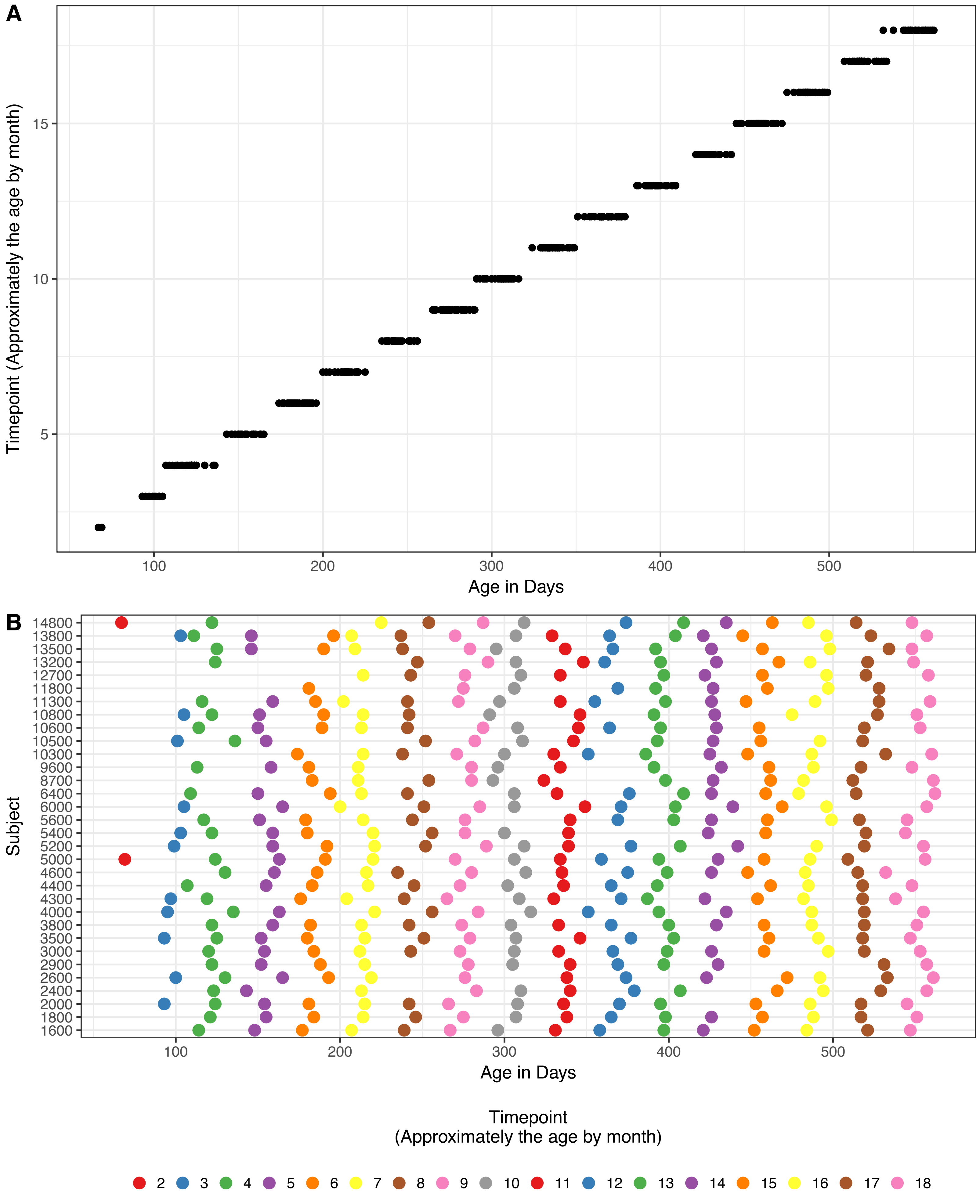
