## Supplementary material for "Interactions between the gut microbiome and mucosal immunoglobulins A, M and G in the developing infant gut": Figure S11

Supplementary figure S11

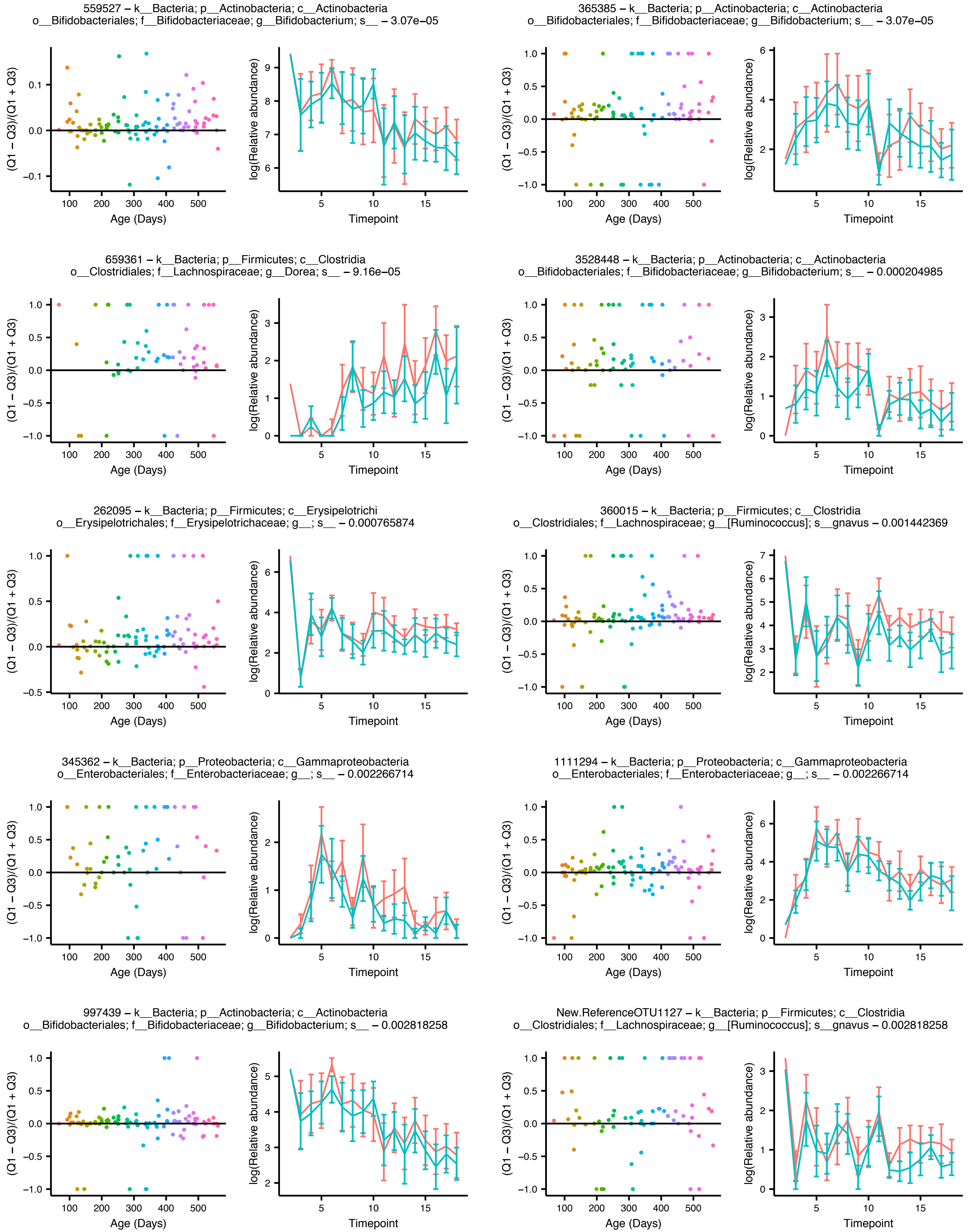

Figure Teddy_Q1_vs_Q3_supp

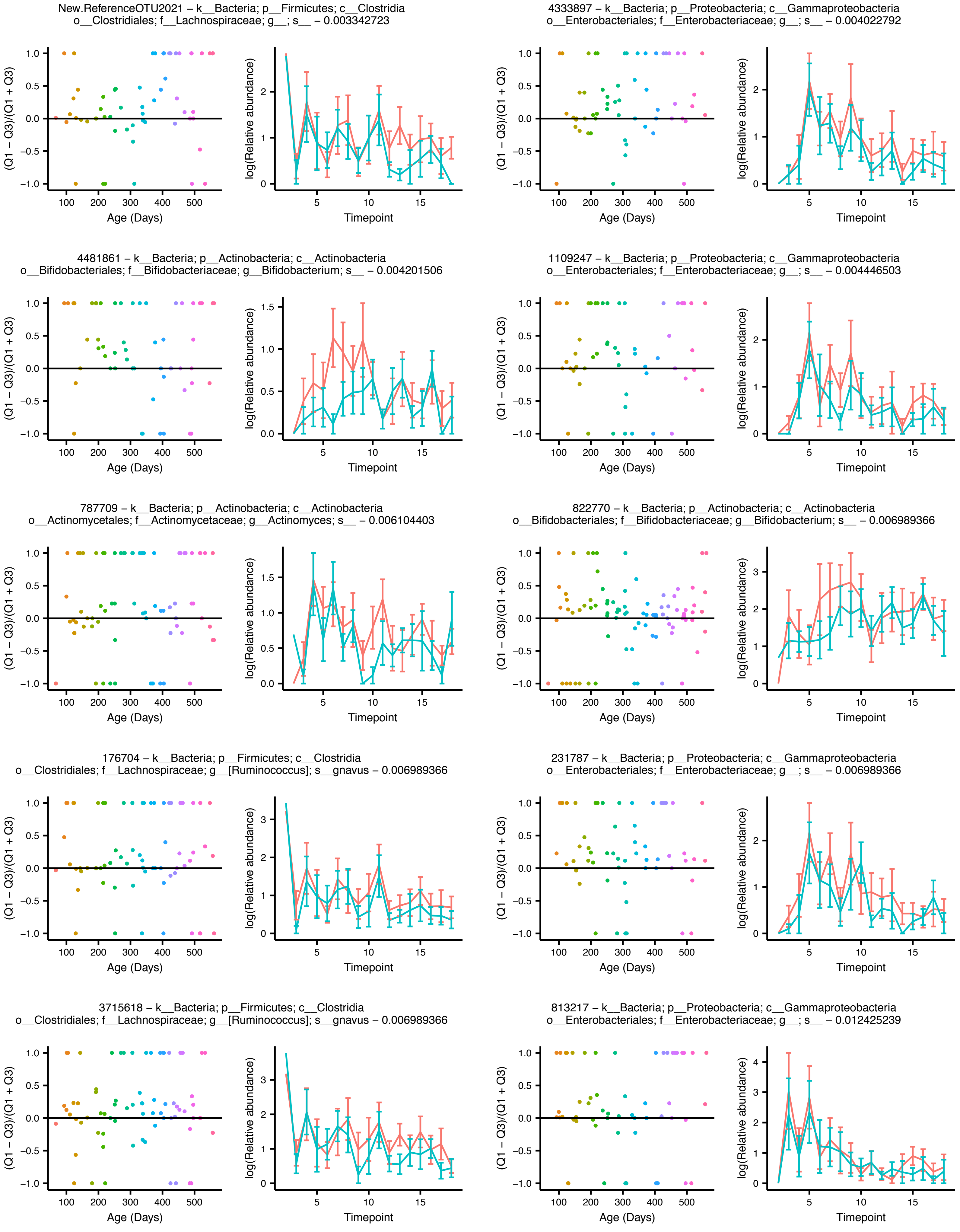

### Figure Teddy_Q1_vs_Q3_supp

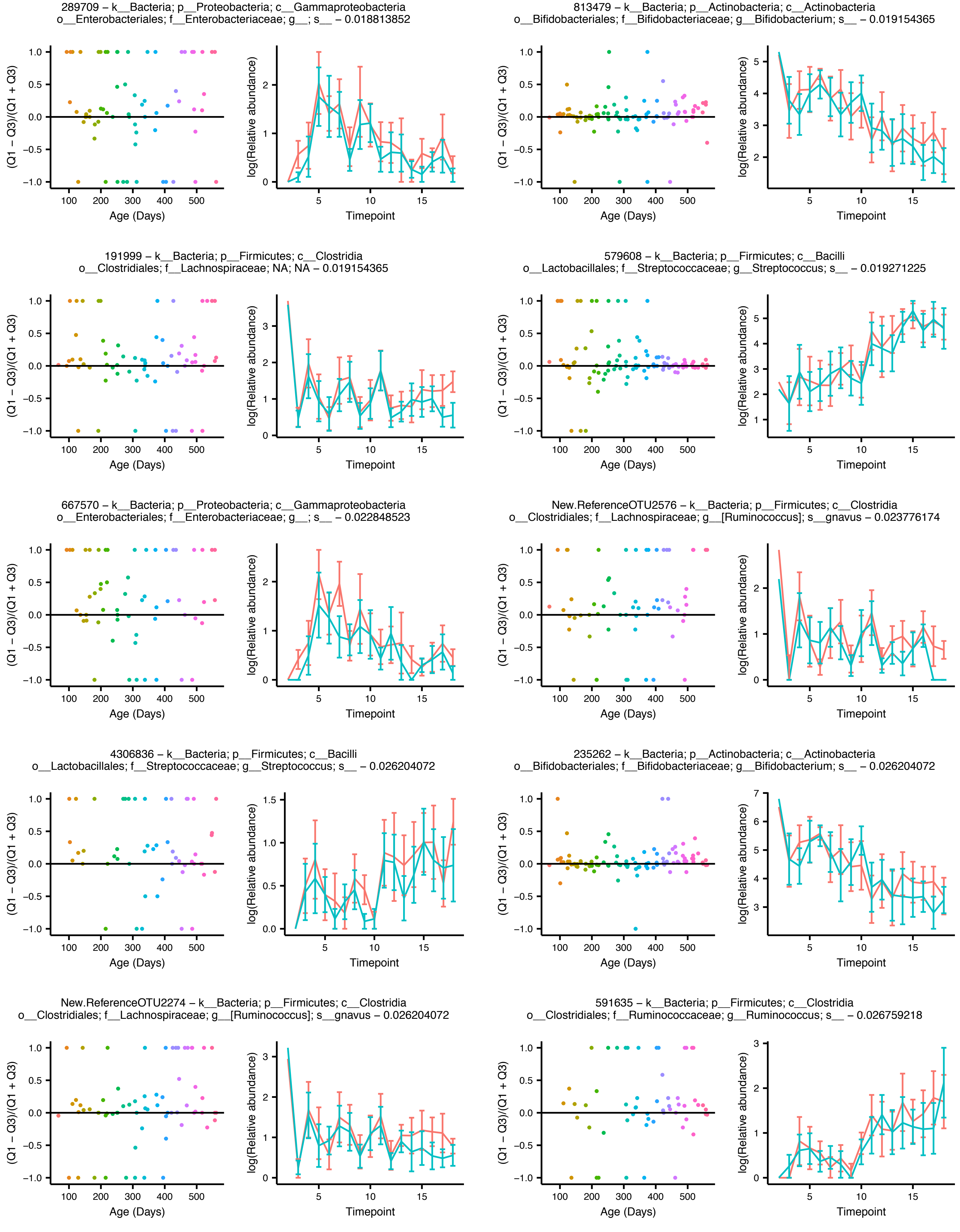

Figure Teddy_Q1_vs_Q3_supp

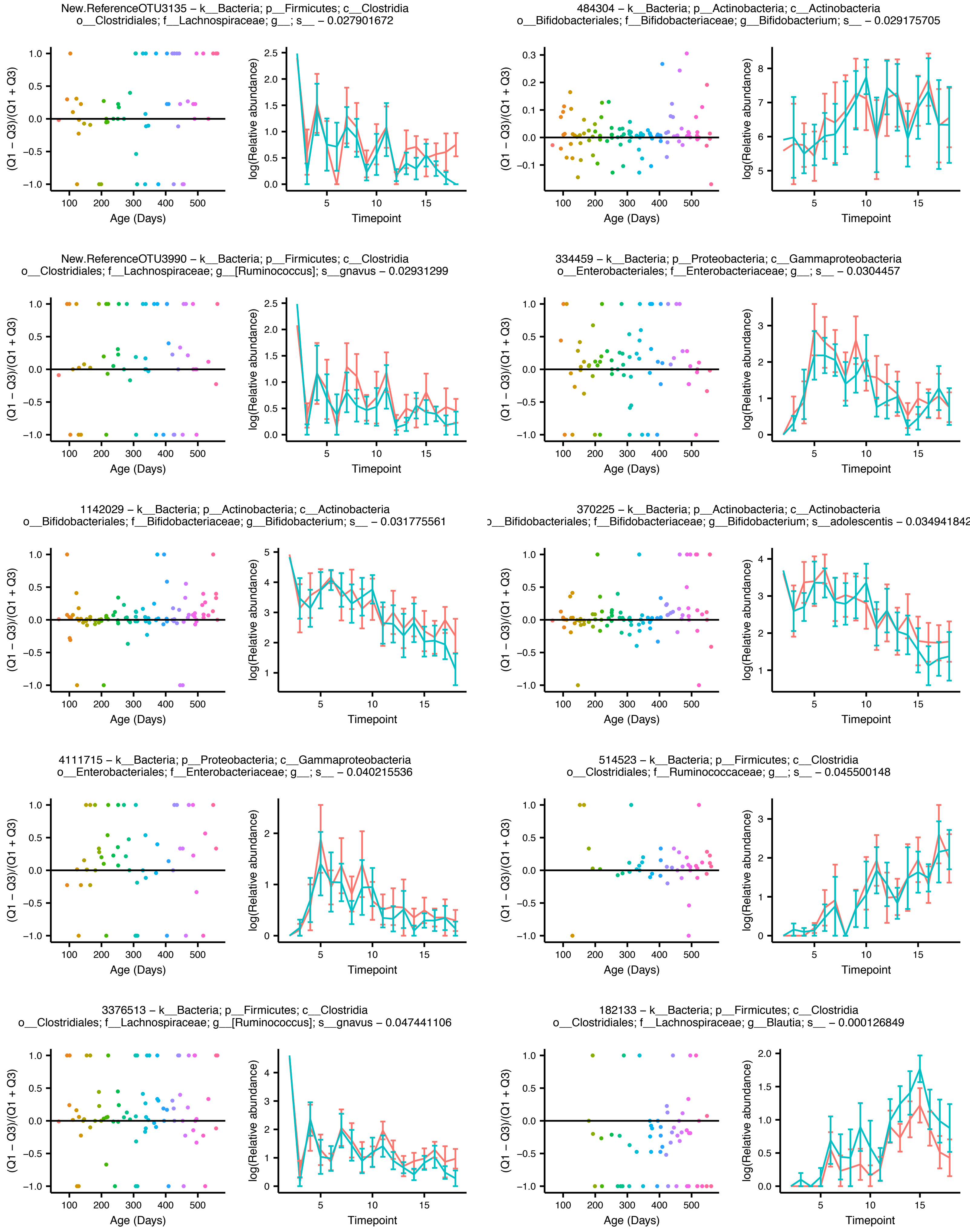

Figure Teddy_Q1_vs_Q3_supp

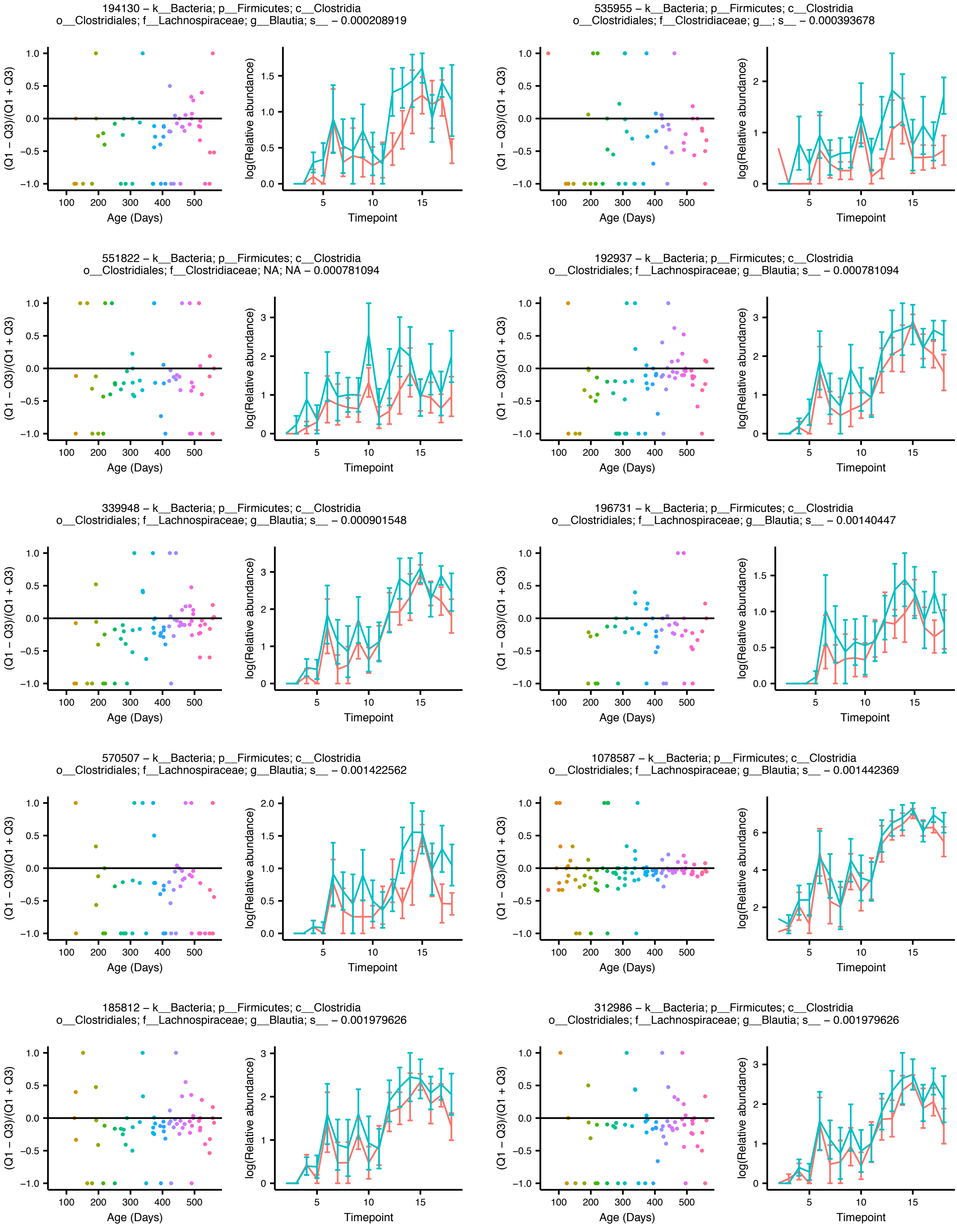

### Figure Teddy_Q1_vs_Q3_supp

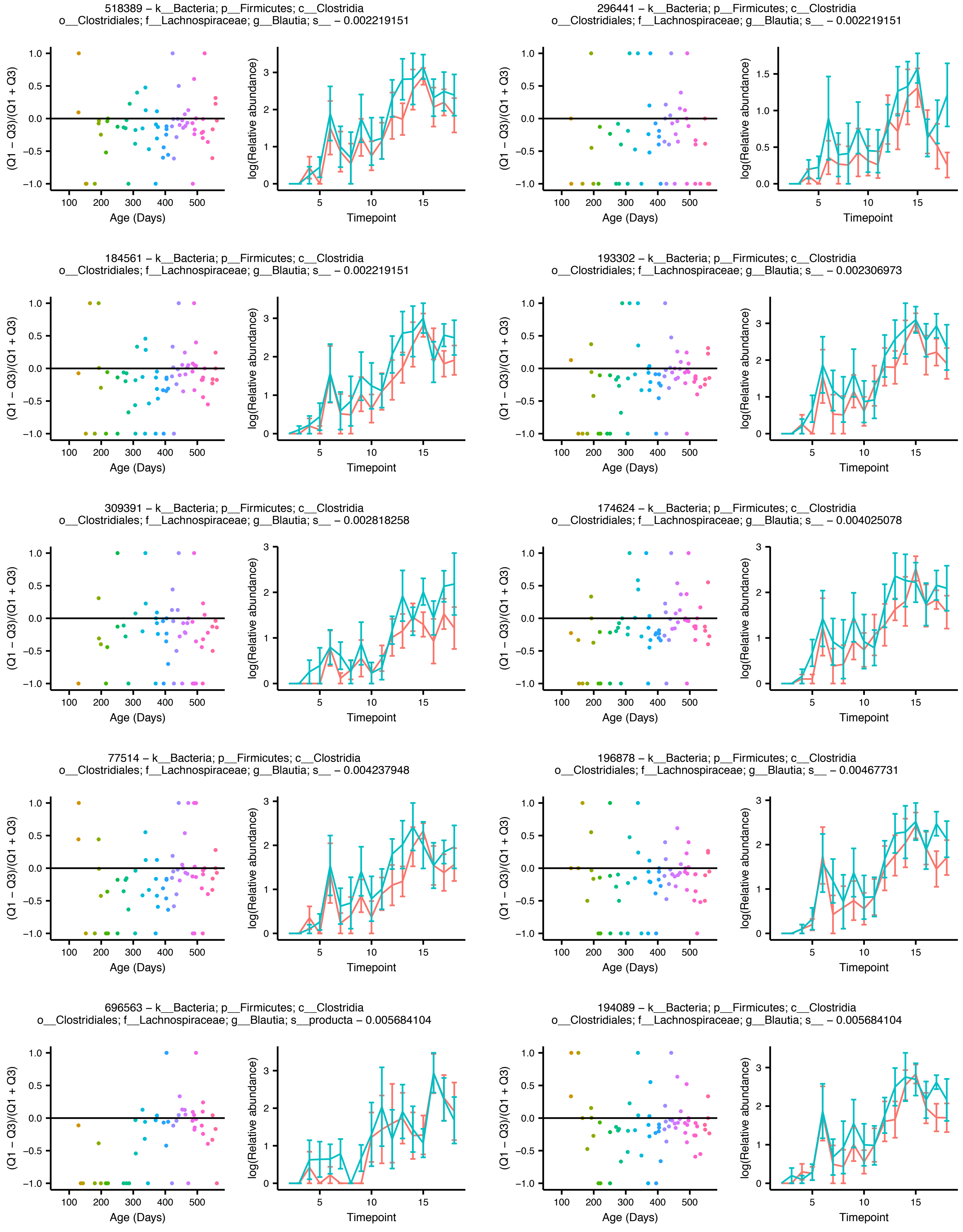

Figure Teddy_Q1_vs_Q3_supp

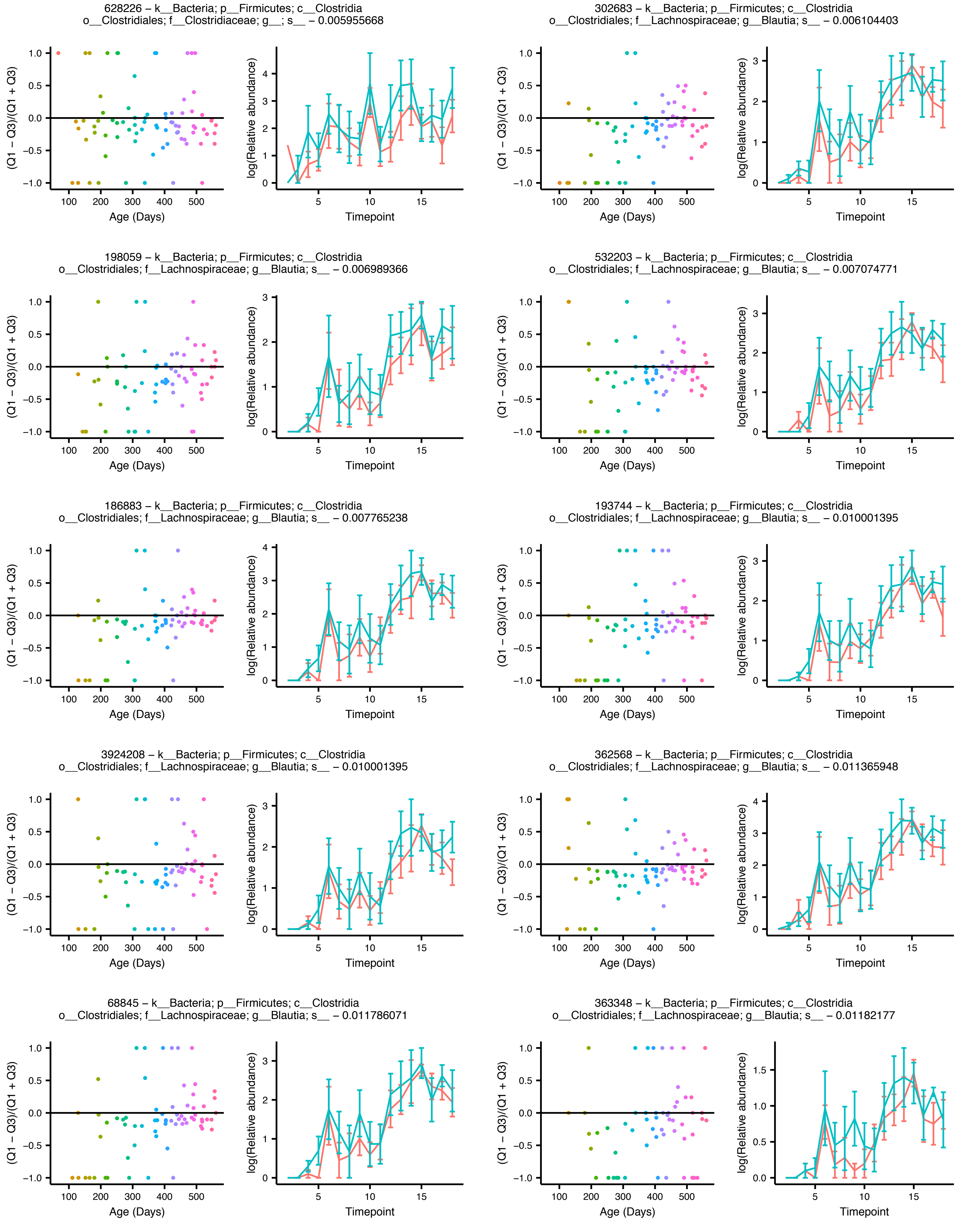

### Figure Teddy_Q1_vs_Q3_supp

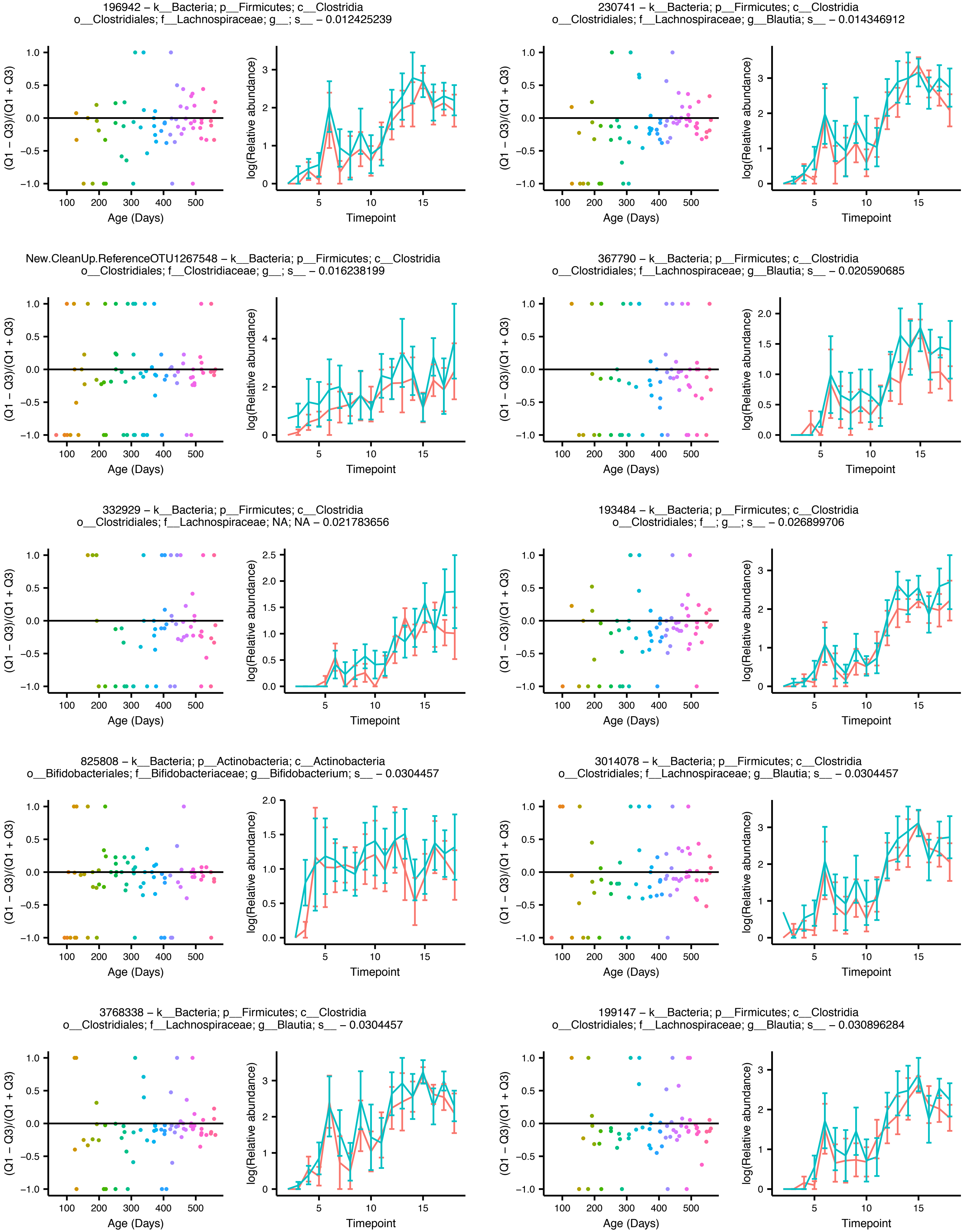
