## Supplementary material for "Interactions between the gut microbiome and mucosal immunoglobulins A, M and G in the developing infant gut": Figure S12

Supplementary figure S12

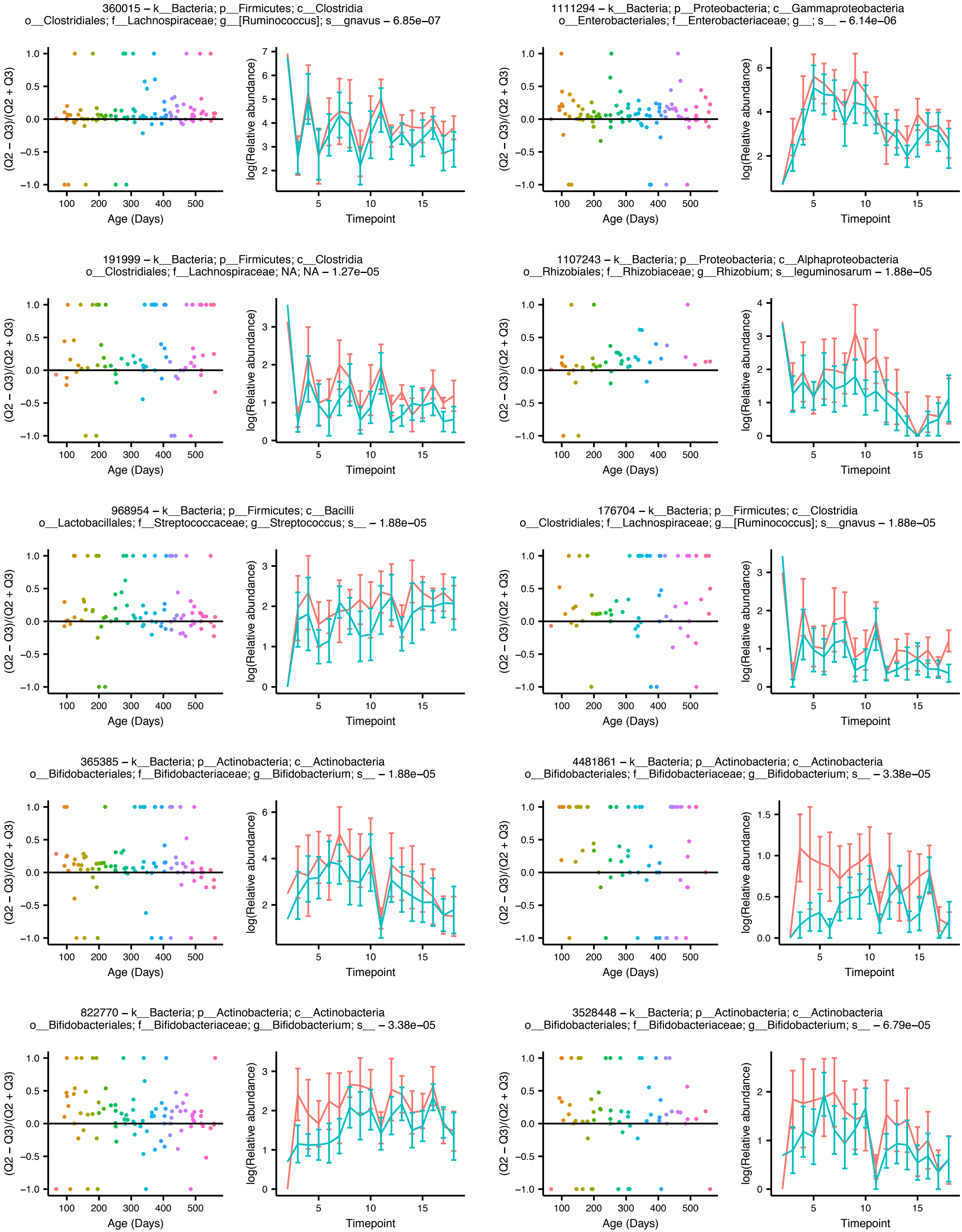

### Figure Teddy_Q2_vs_Q3_supp

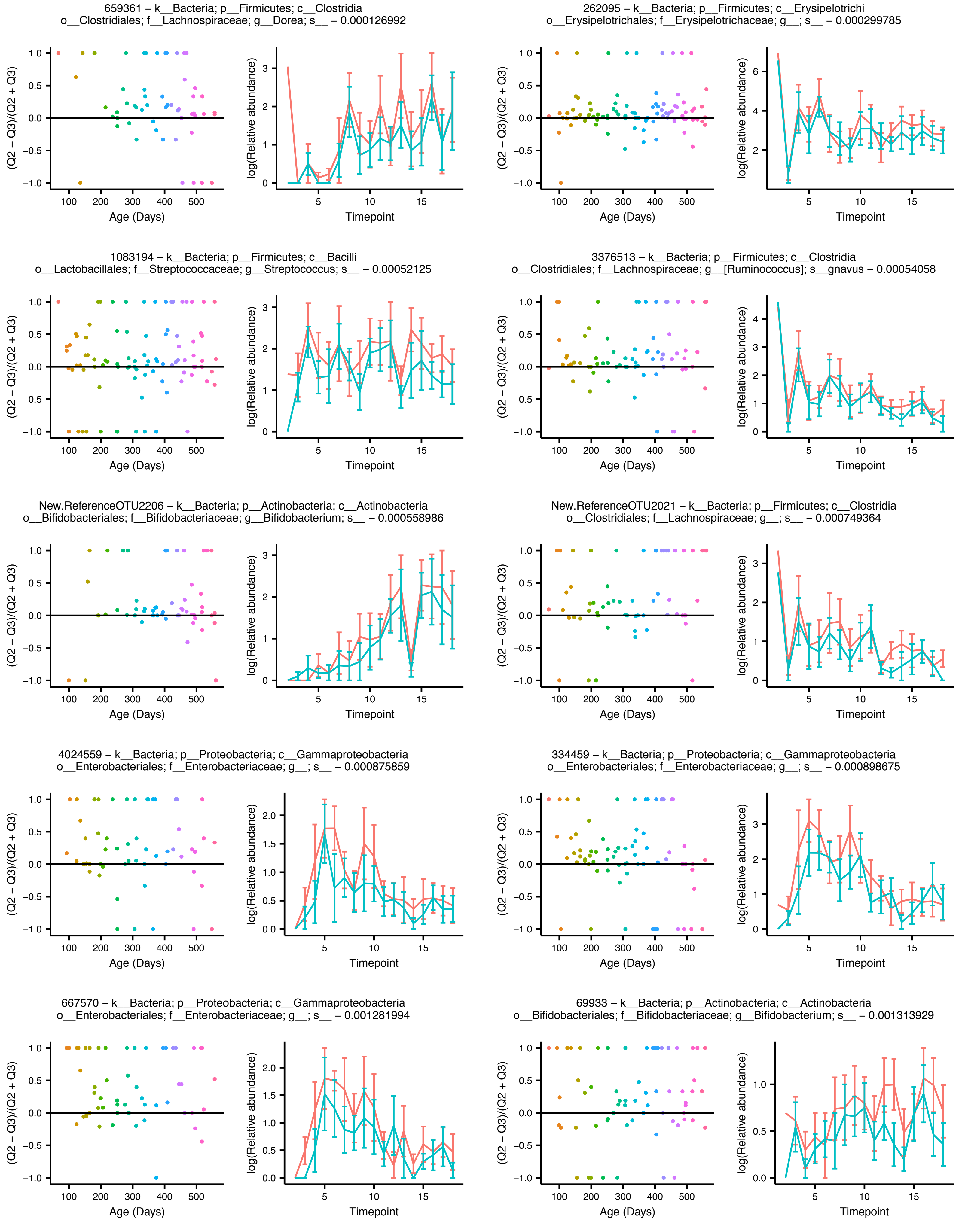

### Figure Teddy_Q2_vs_Q3_supp

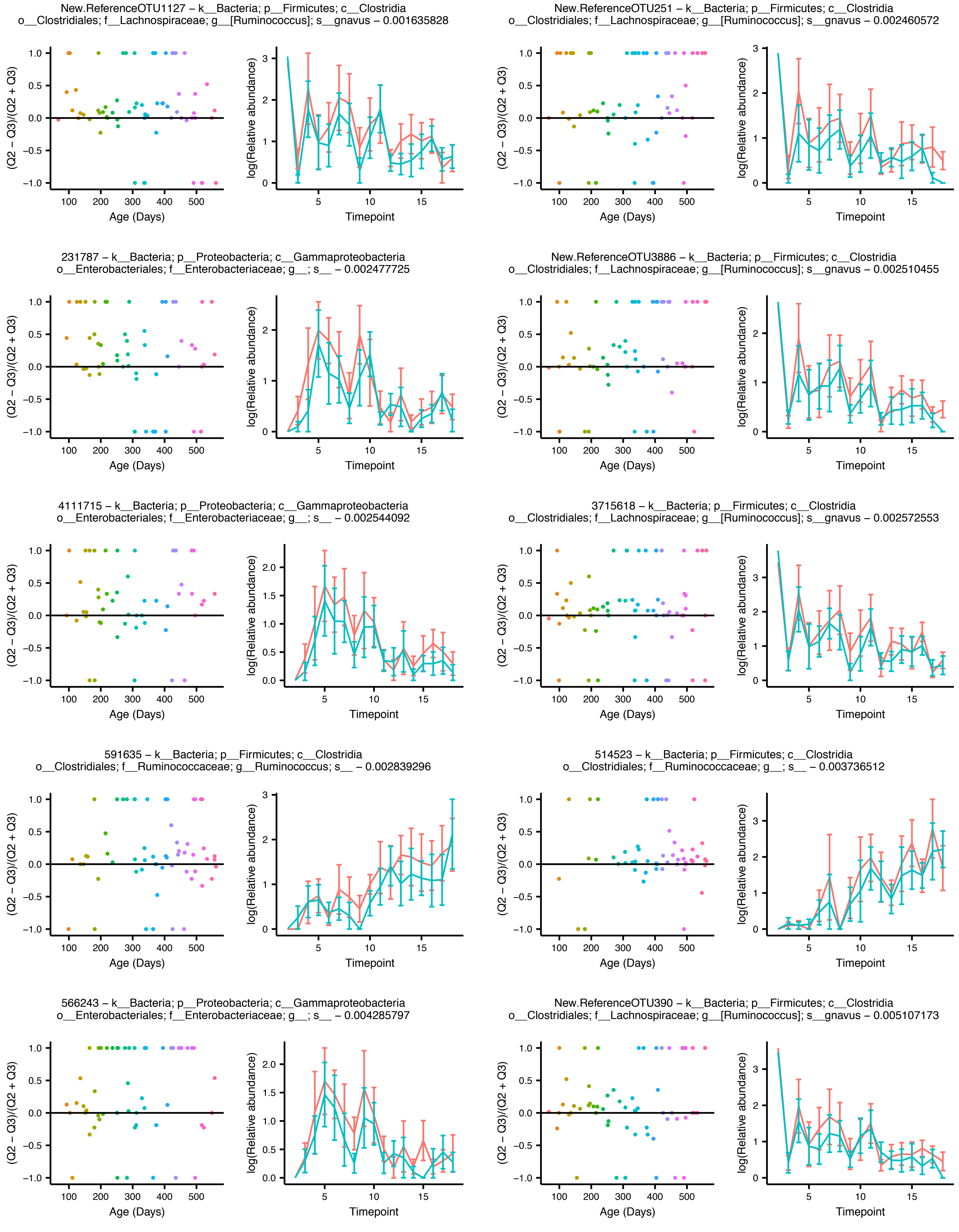

### Figure Teddy_Q2_vs_Q3_supp

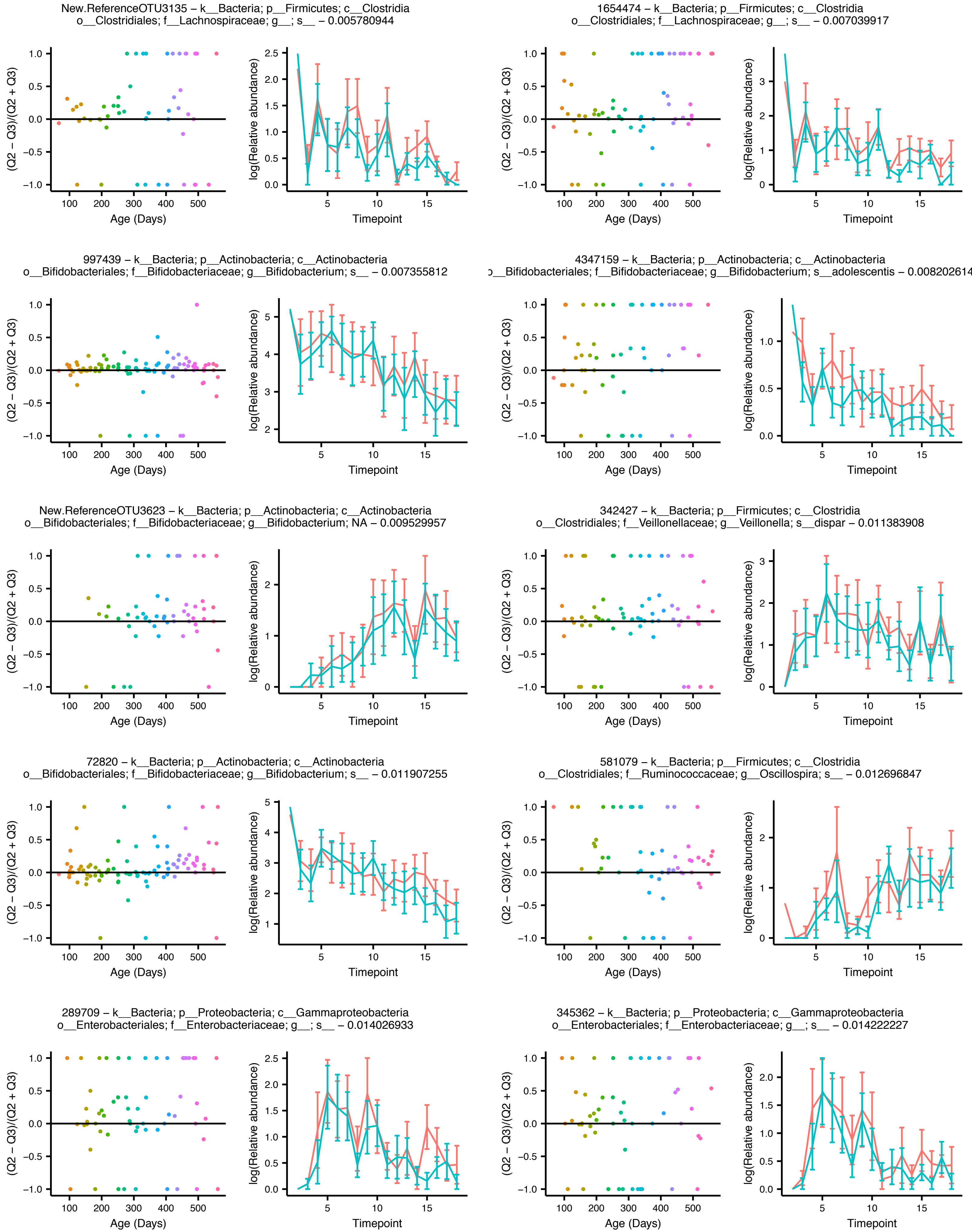

### Figure Teddy_Q2_vs_Q3_supp

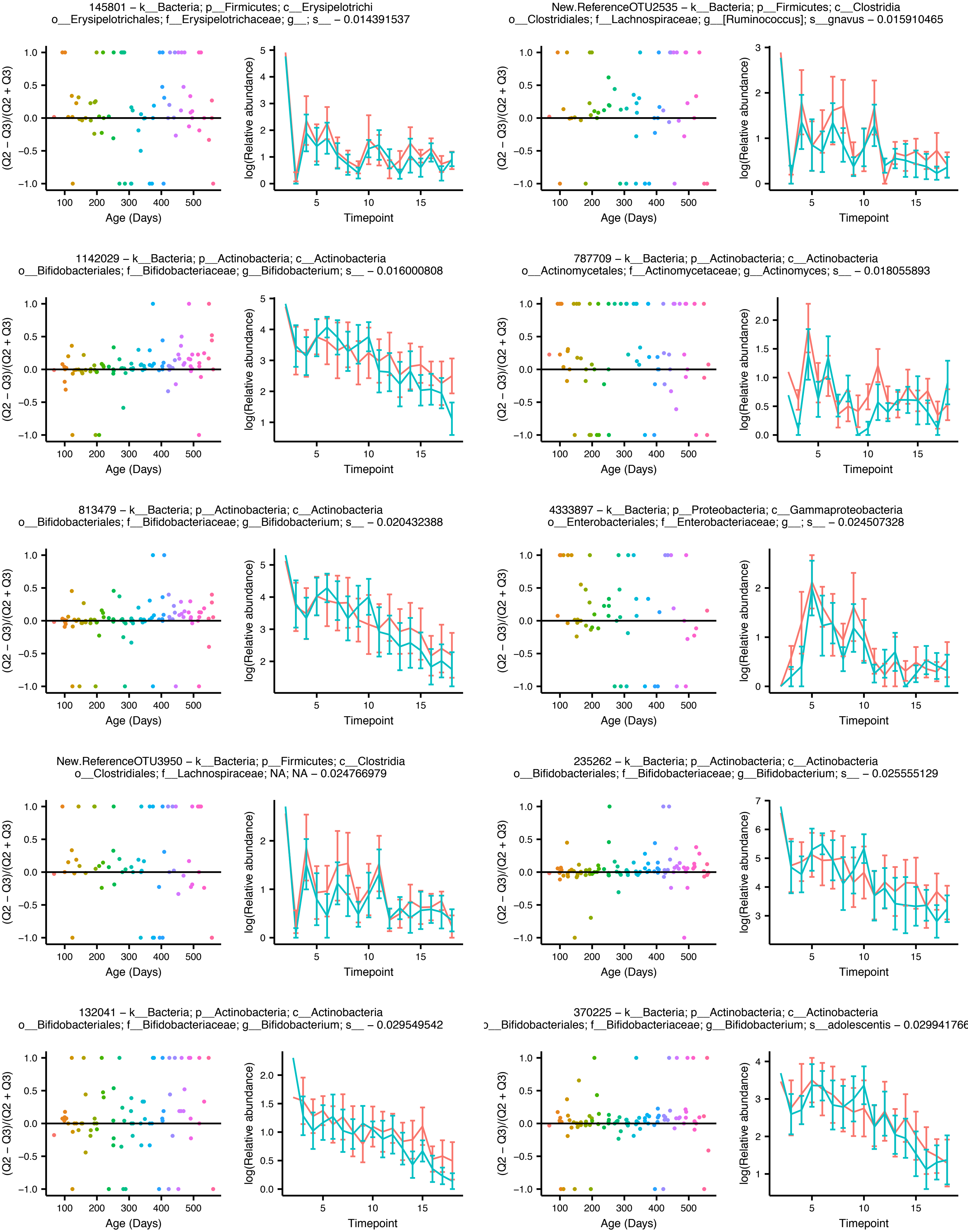

#### Figure Teddy_Q2_vs_Q3_supp

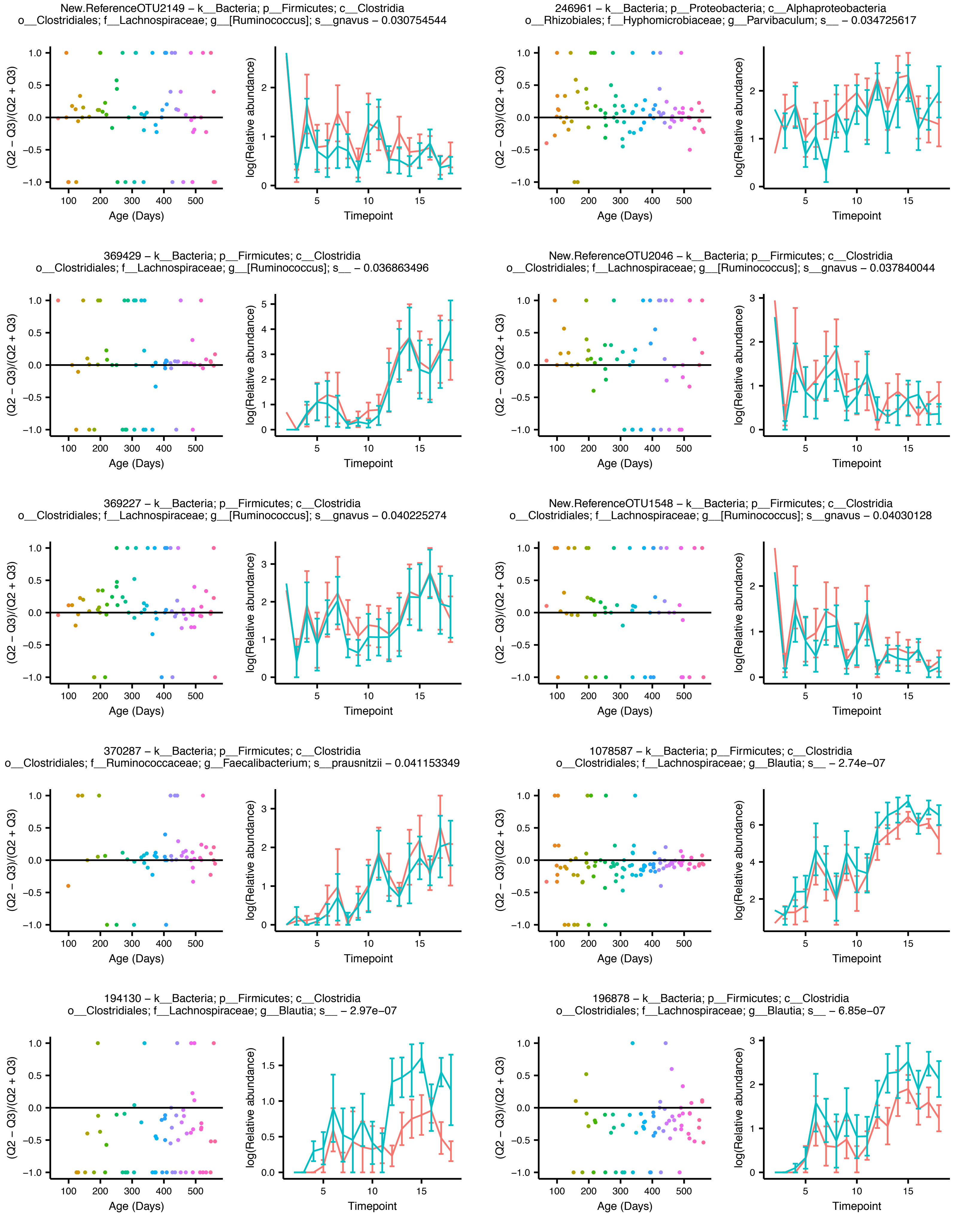

### Figure Teddy_Q2_vs_Q3_supp

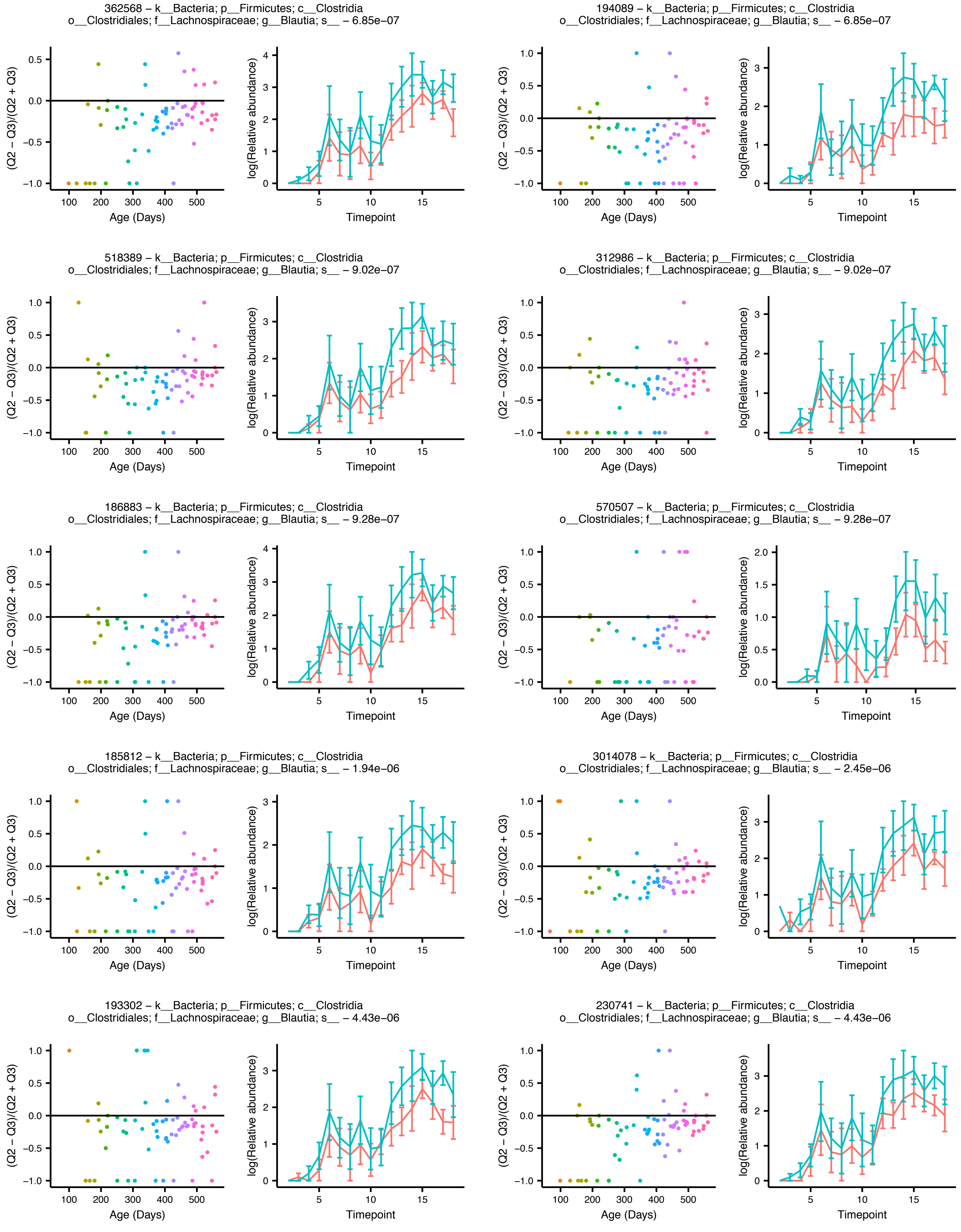

### Figure Teddy_Q2_vs_Q3_supp

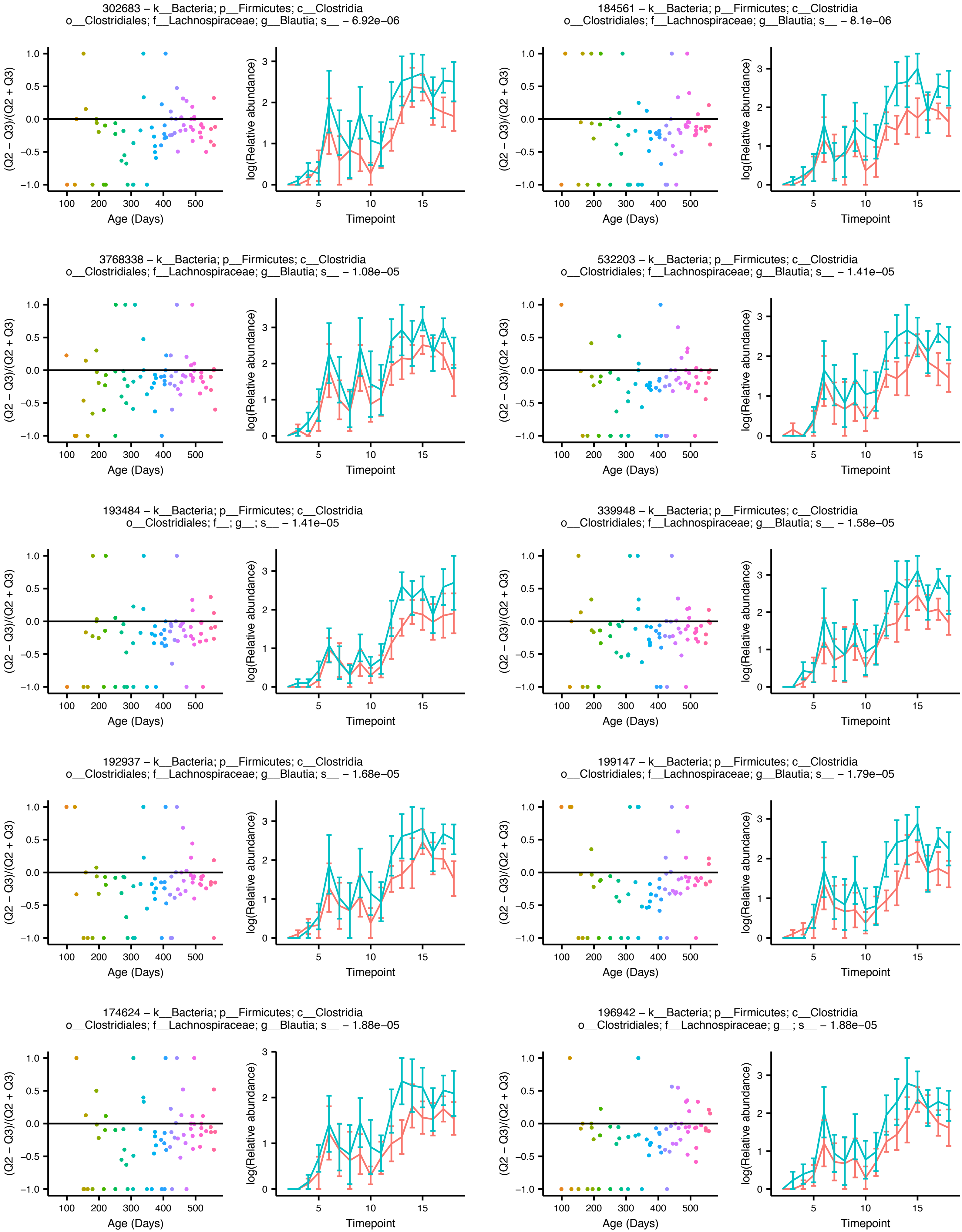

Figure Teddy_Q2_vs_Q3_supp

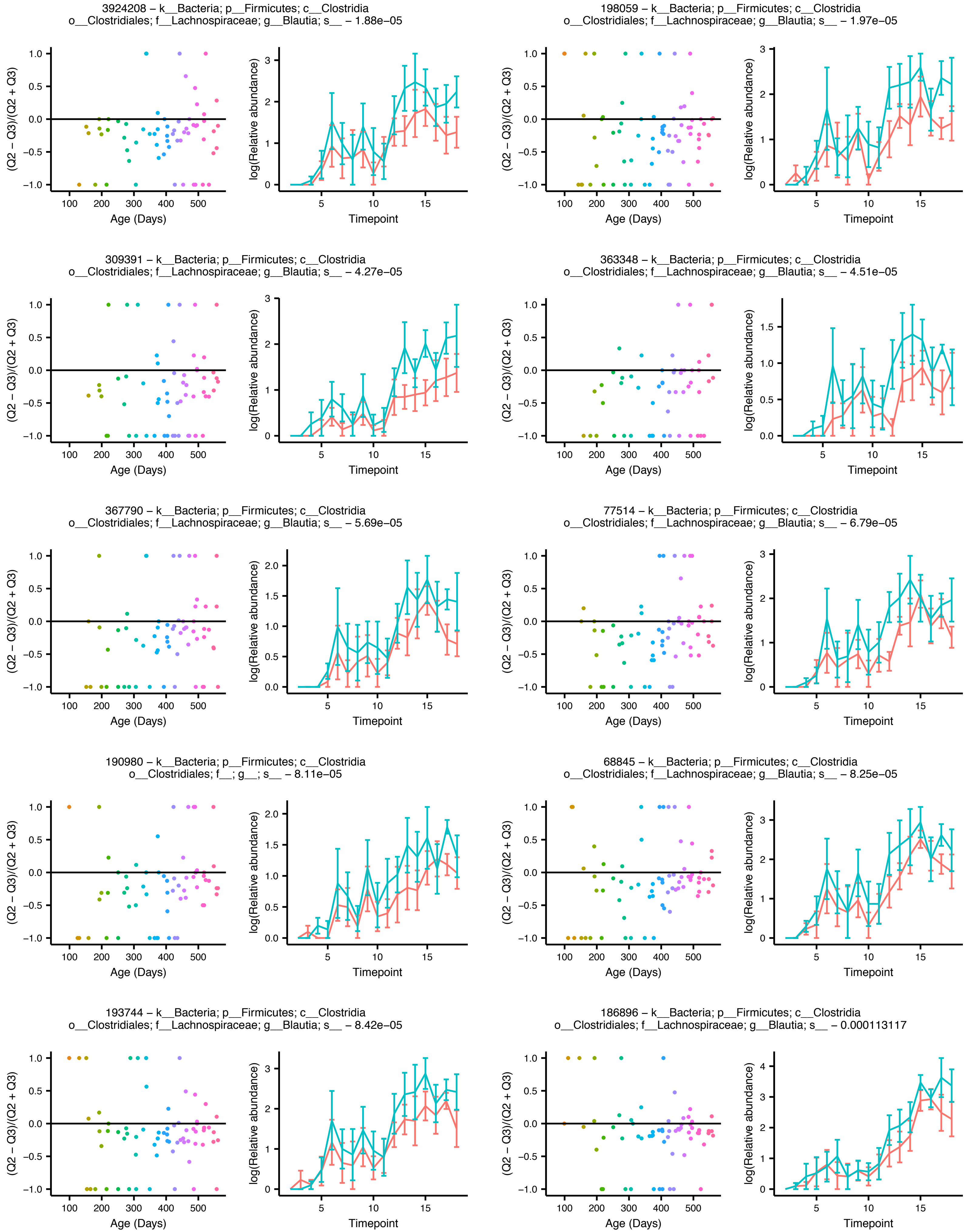

### Figure Teddy_Q2_vs_Q3_supp

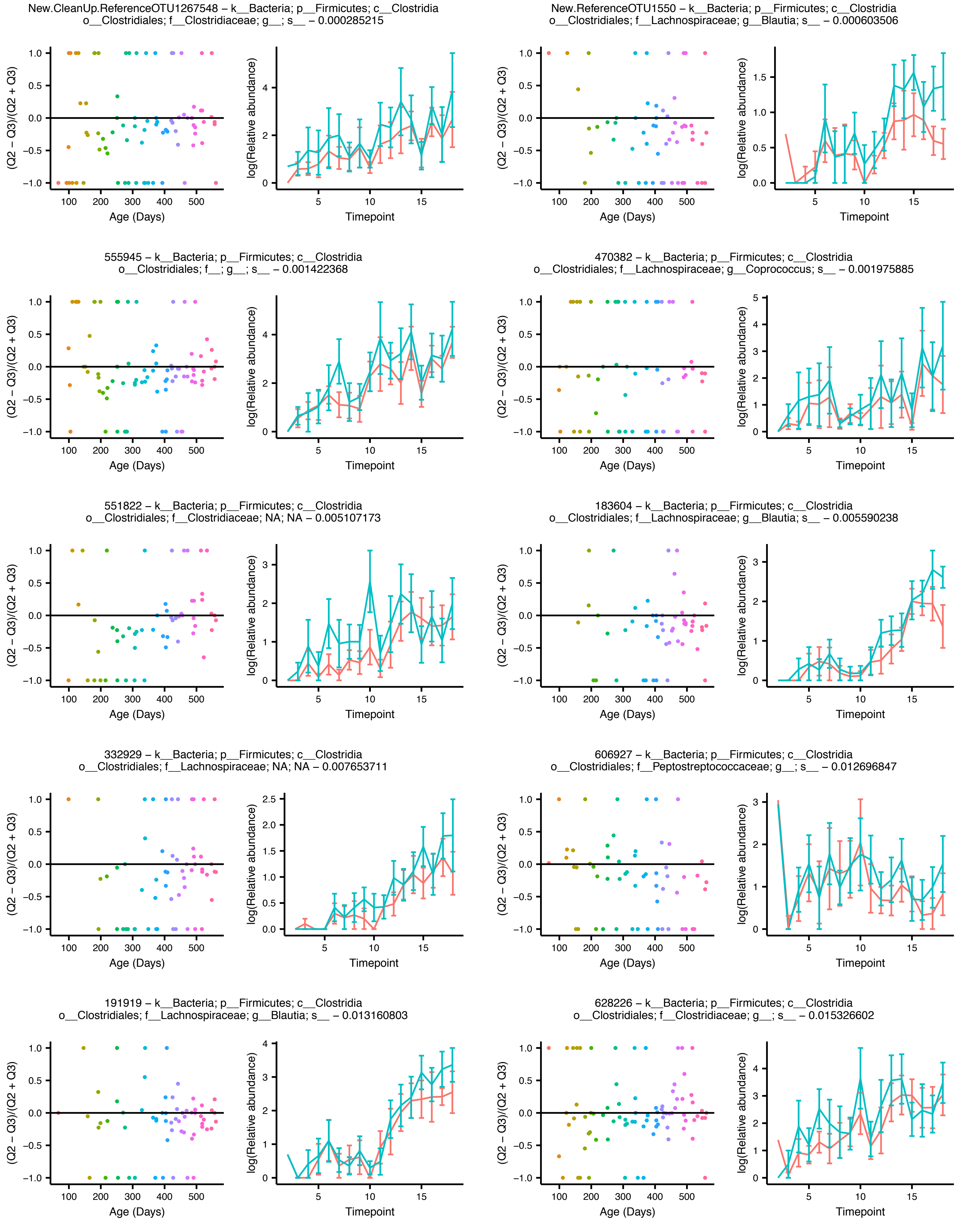

### Figure Teddy_Q2_vs_Q3_supp

New.ReferenceOTU2357 – k__Bacteria; p__Firmicutes; c__Clostridia
o__Clostridiales; f__Lachnospiraceae; g__Blautia; s__ – 0.026606995

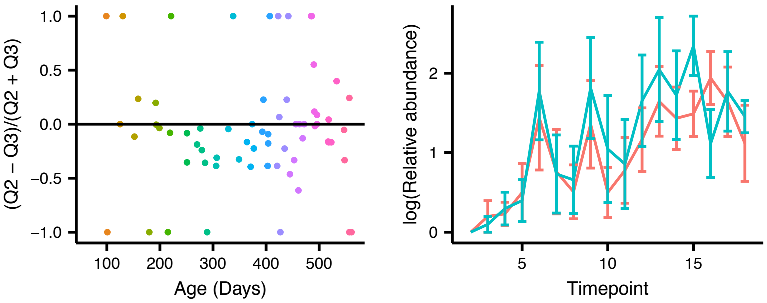
